# Genes adapt to outsmart gene targeting strategies in mutant mouse strains by skipping exons to reinitiate transcription and translation

**DOI:** 10.1101/2020.04.22.041087

**Authors:** Vishnu Hosur, Benjamin E. Low, Daniel Li, Grace A. Stafford, Vivek Kohar, Leonard D. Shultz, Michael V. Wiles

**Affiliations:** The Jackson Laboratory for Mammalian Genetics, Bar Harbor, ME.; Arthritis and Tissue Degeneration Program, Hospital for Special Surgery at Weill Cornell Medicine, New York, NY, 10021

## Abstract

Gene disruption in mouse embryonic stem cells or zygotes is a conventional genetics approach to identify gene function *in vivo*. However, because different gene-disruption strategies use different mechanisms to disrupt genes, the strategies can result in diverse phenotypes in the resulting mouse model. To determine whether different gene-disruption strategies affect the phenotype of resulting mutant mice, we characterized *Rhbdf1* mouse mutant strains generated by three commonly used strategies—definitive-null, targeted knockout (KO)-first, and CRISPR/Cas9. We find that *Rhbdf1* responds differently to distinct KO strategies, for example, by skipping exons and reinitiating translation to potentially yield gain-of-function alleles rather than the expected null or severe hypomorphic alleles. Our analysis also revealed that at least 4% of mice generated using the KO-first strategy show conflicting phenotypes, suggesting that exon skipping is a widespread phenomenon occurring across the genome. Additionally, our study emphasizes that at least 35% of mouse and 45% of human protein-coding genes could be predisposed to targeted KO-first- and CRISPR/Cas9-mediated unexpected translation. Our findings have significant implications for the application of genome editing in both basic research and clinical practice.

## Introduction

Mice are closely related, genetically, to humans, and hence mice have been chosen as a model system to decipher the function of ~20,000 protein-coding genes to gain insights into human biology and disease. For large-scale mouse mutagenesis efforts, gene targeting by homologous recombination in mouse embryonic stem (ES) cells was an efficient and versatile technique. Gene targeting involved either a definitive-null design, which deleted the entire genomic sequence of the target gene, or a targeted knockout (KO)-first design, which offered several advantages, including a gene disruption and a reporter-tagged mutation, and furthermore, permitted analysis of gene function in a tissue-specific or temporal manner. Recently, the use of CRISPR/Cas9 to disrupt genes directly in zygotes has superseded both the definitive-null and KO-first strategies. To determine whether different gene-targeting-strategies affect the phenotype of homozygous mutant mice, we systematically characterize *Rhbdf1* mutant mice generated by these three KO strategies—definitive-null, targeted KO-first, and CRISPR/Cas9. The *Rhbdf1* gene encodes RHBDF1 and has been implicated as playing a key role in growth and development [1], inflammation [2] and cancer [3–5].

Definitive-null and targeted KO-first strategies are powerful high-throughput approaches that can be used for large-scale gene targeting in ES cells to study thousands of mammalian protein-coding genes to better understand human biology and disease [6–8]. In the use of the definitive-null strategy, bacterial artificial chromosome (BAC)-based targeting vectors replace the entire genomic sequence of the target gene (Supplementary Fig. 1a), thereby generating a null allele. In contrast, the targeted KO-first approach [9, 10], a strategy that includes alternative steps that can be chosen based on the desired result, is highly versatile, as it enables the analysis of gene function in a tissue-specific or temporal manner. The KO-first allele is created through targeted delivery of a cassette that includes a mouse *En2* splice site, a β-galactosidase reporter, a neomycin resistance gene, and a transcriptional termination SV40 polyadenylation sequence, into the intronic sequence resulting in the termination of transcription of the targeted gene is terminated by the polyadenylation sequence before the entire gene sequence is transcribed, resulting in truncated transcription of a nonfunctional reporter protein. Application of cre recombinase to a KO-first ES cell clone results in generation of a null allele (Supplementary Fig. 1b). Application of Flp recombinase to a KO-first ES cell clone converts the KO-first allele into a conditional-ready (or simply a wildtype) allele by removing the cassette flanked by the *FRT* sites (Supplementary Fig. 1c); subsequent application of cre recombinase elegantly generates conditional alleles by deleting critical exon(s) that are flanked by *loxP* sites (Supplementary Fig. 1c). The “KO-first” alleles, by offering substantial flexibility with respect to either gene disruption or target tissue expression and timing, are time-efficient. Despite evidence suggesting that targeted KO-first alleles may result in hypomorphs [11], to our knowledge there have been no reports indicating that use of the targeted KO-first strategy has led to gain-of-function alleles that rescue the severe phenotype caused by the null alleles.

CRISPR/Cas9-mediated genome editing is currently the primary conventional approach for generating null alleles [12]. This method relies on the introduction of frameshift mutations via non-homologous end joining (NHEJ) in ES cells or zygotes to trigger nonsense-mediated decay (NMD) of mRNA transcripts [13]. However, emerging evidence suggests that these mutant alleles can reinitiate translation from a downstream start codon resulting in truncated proteins [14]. Recently, two separate studies performed systematic analysis of CRISPR/Cas9-induced frameshift mutations in HAP1 cells and revealed that one third of the genes could generate truncated proteins either by reinitiating translation or skipping the edited exon [15, 16]. Likewise, in zebrafish, CRISPR/Cas9-mediated generation of KO alleles resulted in unpredicted mRNA transcripts owing to exon skipping, nonsense-associated altered splicing, or use of cryptic splice sites [17]. Collectively, these studies highlight the limitations of CRISPR/Cas9-mediated generation of null alleles.

Building on the aforementioned work in HAP1 cell lines and mutant zebrafish lines, we comprehensively characterized *Rhbdf1* mutant mouse strains generated by three distinct gene targeting strategies—definitive-null, targeted KO-first, and CRISPR/Cas9—using whole exome sequencing (Exome-Seq), RNA sequencing (RNA-Seq), 5’-rapid amplification of cDNA ends (RACE), and functional analysis of residual proteins. We find that whereas definitive-null *Rhbdf1* mutant mice (hereafter referred to as *Rhbdf1*^*null/null*^) show growth retardation, brain and heart defects, and the death of the majority of mice by postnatal day 14 (P14), targeted KO-first- and CRISPR/Cas9-engineered *Rhbdf1* mutant mice do not display growth retardation or any of the other pathologies observed in the *Rhbdf1*^*null/null*^ mice, suggesting incomplete elimination of RHBDF1 biological activity. Comprehensive analyses of all three *Rhbdf1* mutant alleles revealed the existence of functional N-terminally truncated RHBDF1 mutant proteins capable of rescuing the multi-organ pathology observed in *Rhbdf1*^*null/null*^ mice. Our data suggest that because mutant transcripts are capable of generating functional truncated proteins, a thorough post-genomic-DNA analysis of targeted KO-first- and CRISPR/Cas9-engineered alleles, and additional studies as well, are warranted to validate the null alleles before phenotypic characterization of mutant mouse strains.

## Results

### Does the choice of gene targeting strategy affect the phenotype?

In recent studies, two different groups each generated *Rhbdf1* homozygous mutant mice, and reported conflicting results regarding the phenotypes [1, 18]; Christova and colleagues, using the definitive-null strategy, generated an *Rhbdf1* loss-of-function mutation in two different inbred mouse strains: C57BL/6J and 129S6. In both strains, heterozygous-null mice were normal and indistinguishable from wildtype littermates, whereas homozygous-null mice had low body weights and died at about 9 days (C57BL/6J) and 6 weeks (129S6) after birth (Fig. 1a, see A1). In contrast, using the targeted KO-first strategy, Li and colleagues reported that *Rhbdf1* mutant mice generated on a mixed genetic background—C57BL/6N and FVB—were healthy with no apparent phenotypic or histological abnormalities, and proposed that RHBDF1 is not essential for mouse development (Fig. 1a, see A3). Furthermore, Christova et al. (using both C57BL/6J and 129S6 mice) and Li et al. (using mice on the mixed C57BL/6N and FVB genetic background) also both generated *Rhbdf1:Rhbdf2* double mutant mice, and the phenotypes differed considerably between the two studies. While Christova et al. reported *in utero* lethality in the double mutants (Fig. 1a, see A2), Li et al. observed perinatal lethality with an ‘eyelids open at birth’ phenotype and heart-valve defects (Fig. 1a, see A4). It seems unlikely that there is a modifier in the FVB background that could account for the difference in the phenotype observed in the single and double mutant mice generated on the C57BL/6J background by Christova et al. It seems more likely, rather, that the differences in the phenotypes could be due to effects of the different gene targeting strategies employed (Fig. 1a) by the two studies to generate *Rhbdf1* KO mice.

**Figure 1.**
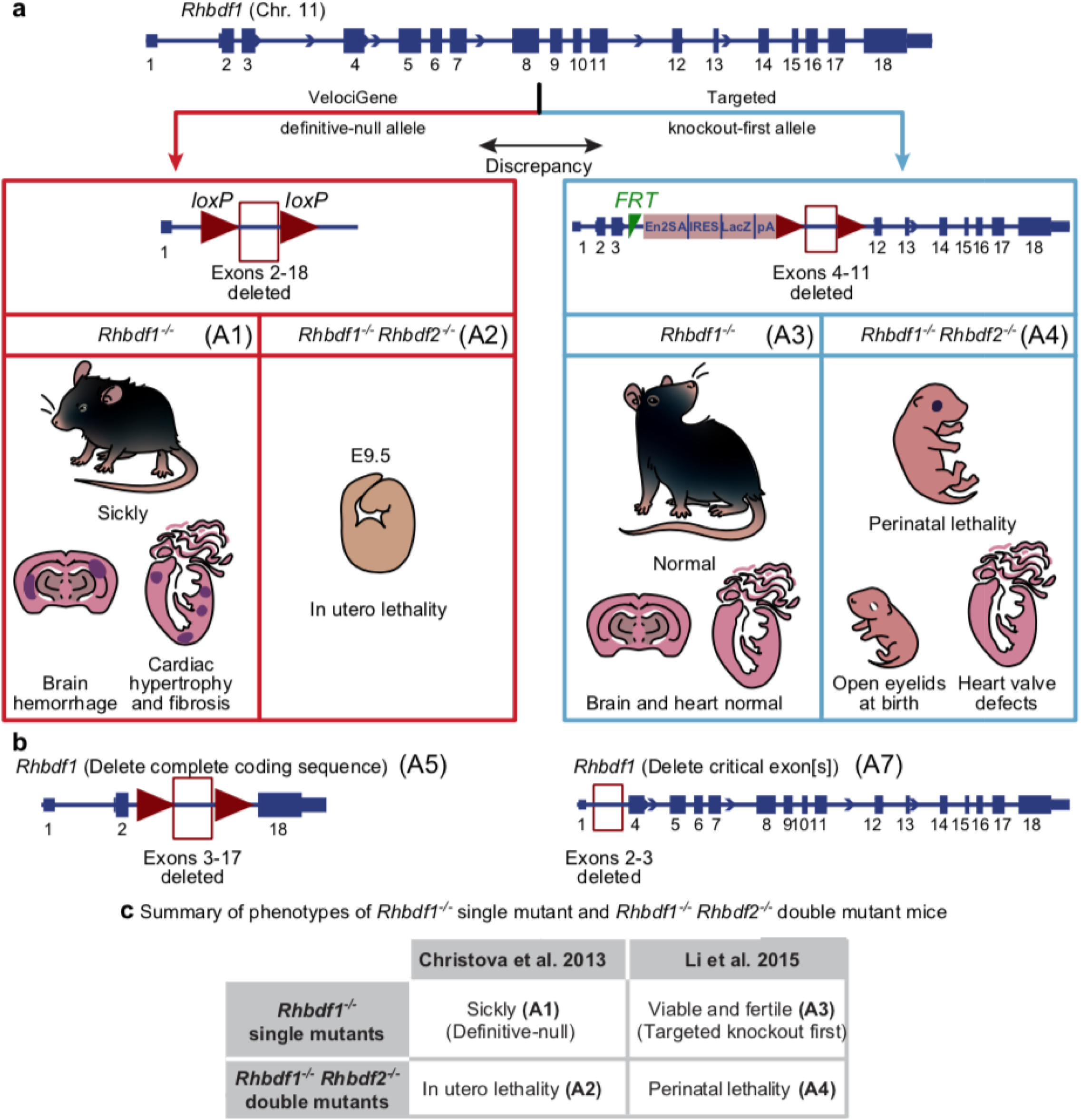
Analysis of *Rhbdf1*-deficient mice suggests that the targeting strategy might affect the phenotype of homozygous mutant mice. a. Schematic representation of exons (filled boxes) and introns (lines) in the *Rhbdf1* gene. Previous studies have produced conflicting results regarding the consequences of *Rhbdf1* inactivation in mice. Christova et al. reported that *Rhbdf1*^*−/−*^ mice generated using the VelociGene definitive-null gene KO strategy suffer from multi-organ pathologies and die within ~6 weeks (A1), and that the *Rhbdf1*^*−/−*^ *Rhbdf2*^*−/−*^ (A2) double KO mice, generated using A1 *Rhbdf1*^*−/−*^ mice, died prenatally with phenotypes that were significantly more severe. In contrast, Li et al. reported that *Rhbdf1*^*−/−*^ homozygous mutant mice generated ‘with a targeted KO-first allele showed no evidence of pathological phenotypes (A3). Moreover, *Rhbdf1*^*−/−*^ *Rhbdf2*^*−/−*^ double mutant mice, generated using A3 *Rhbdf1*^*−/−*^ mice, exhibited less severe disease phenotypes, such as heart valve defects and an “open eyelids at birth’ phenotype (A4). b. We hypothesized that the targeted KO-first allele (A3), but not the definitive-null allele, generates an N-terminally truncated RHBDF1 protein product from the mutant mRNA that is not degraded by the NMD mechanism, to rescue the overt phenotype observed in RHBDF1-deficient mice (A1). To test our hypothesis, we independently generated *Rhbdf1* homozygous mutant mice using two different strategies: 1) delete complete coding sequence (A5), and 2) delete critical exon(s) (A7). We postulated that whereas the A5 allele would be analogous to the A1 definitive-null allele, the A7 allele would generate a functional truncated mutant RHBDF1 protein, analogous to the targeted KO-first mutant allele A3, rescuing the multi-organ pathology of RHBDF1-deficient mice (A1). c. Table indicating the phenotypes of previously generated *Rhbdf1* single mutant and *Rhbdf1:Rhbdf2* double mutant mice.

To address the underlying cause of discrepancy, we generated a) *Rhbdf1* KO mice using the definitive-null design, which results in alleles that delete the entire protein coding sequence of the target gene (see A5), and b) *Rhbdf1* KO mice using CRISPR/Cas9-mediated gene editing (Fig. 1b, see A7), which results in alleles lacking the critical exon(s) with the translation initiation site ATG (Fig. 1b). We hypothesized that the mutant transcripts generated from the targeted KO-first or CRISPR/Cas9 alleles reinitiate transcription and translation in unexpected ways to rescue the more severe phenotype described by Christova et al. We sought to test our hypothesis by systematically characterizing the *Rhbdf1* single mutant mice (Fig. 1a, see A5 and A7) and *Rhbdf1:Rhbdf2* double mutant mice (Fig. 1a, see A6 and A8) generated in the present study, comparing the phenotypes with those reported in previously published studies (Fig. 1c).

### Complete loss of exons in *Rhbdf1* (A5 allele) results in growth retardation, brain hemorrhage, and cardiac hypertrophy

To explore the function of RHBDF1 *in vivo*, we generated *Rhbdf1* homozygous-null (*Rhbdf1*^*−/−*^) mice from heterozygotes with the A5 allele and validated *Rhbdf1* gene disruption by whole exome sequencing (Exome-Seq) and polymerase chain reaction (PCR). Exome-Seq validated loss of exons 3 through 18, leading to deletion of almost all of the cytosolic N-terminus domain and all of the transmembrane domains (Fig. 2a). Filial crosses of heterozygotes resulted in near-Mendelian ratios, suggesting no embryonic lethality due to RHBDF1 deficiency. Moreover, all *Rhbdf1*^*−/−*^ newborns had a milk spot, and as neonates showed no gross deformities or malformations of major organs, except for growth retardation when compared with *Rhbdf1*^*+/−*^ littermates (Fig. 2c, left). The majority of *Rhbdf1*^*−/−*^ mice were found dead by P14, while the few remaining *Rhbdf1*^*−/−*^ neonates survived only to 3 to 4 weeks (Fig. 2d). These survivors appeared severely dehydrated and lethargic, and both female and male *Rhbdf1*^*−/−*^ survivors exhibited significantly lower body weight compared with *Rhbdf1*^*+/−*^ littermates (Fig. 2e).

**Figure 2.**
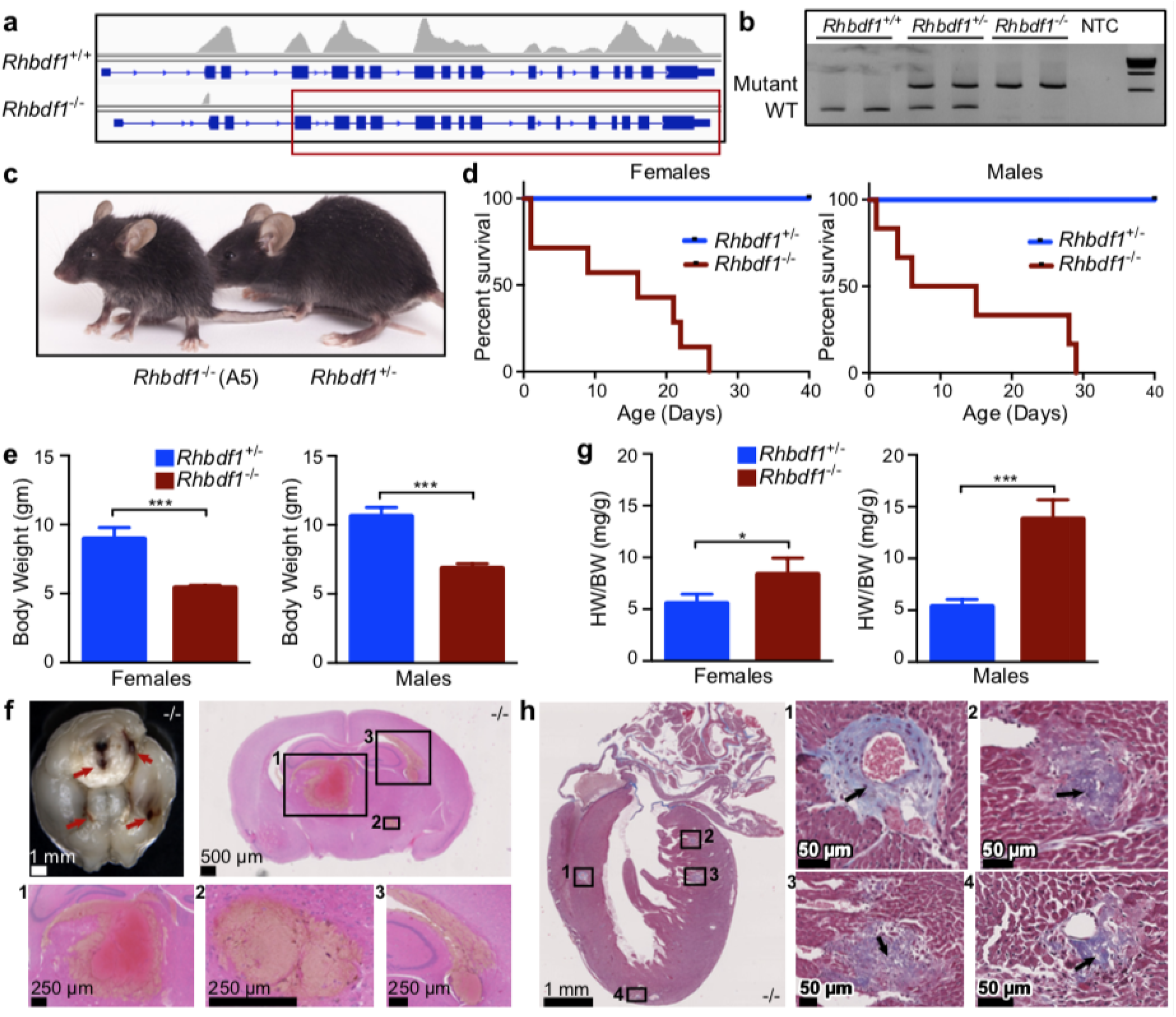
Complete deletion of exons in *Rhbdf1* results in growth retardation, brain hemorrhage, and cardiac hypertrophy (A5) a. Exome-Seq showing loss of exons 2 through 18 in *Rhbdf1*^*−/−*^ homozygous mutant mice generated using the definitive-null strategy. The red rectangle shows the exons lost in the mutant mice. b. PCR of tail genomic DNA with all four primers shows that wildtype mice have one wildtype band (268-bp PCR product), heterozygous-null mice carry a wildtype band (268 bp) and a mutant band (546 bp), and homozygous-null mice carry one mutant band (546 bp). NTC, no template control. New England Biolabs 1 kb DNA Ladder was used as a molecular marker. Samples were run in duplicate. c. Images of representative postnatal day 28 *Rhbdf1*^*−/−*^ (left) and *Rhbdf1*^*+/−*^ (right) male littermates. d. Kaplan-Meier survival analyses showing that the lifespan of *Rhbdf1*^*−/−*^ homozygous-null female (left panel) and male (right panel) mice is significantly shorter than that of heterozygous-null littermate controls. e. Body weights of *Rhbdf1*^*−/−*^ female (n=4) and male (n=3) mice compared with those of *Rhbdf1*^*+/−*^ littermates (n=3 for each sex) on postnatal day 21. Data represent mean ± S.D; ***p<0.001. f. Brain hemorrhaging in *Rhbdf1*^*−/−*^ mice. Upper left panel: multifocal hemorrhages (arrows) were observed in *Rhbdf1*^*−/−*^ mice. All other panels: H&E-stained coronal sections of postnatal day 21 *Rhbdf1*^*−/−*^ brain showing prominent hemorrhages in the brain stem, hippocampus, and hypothalamus. g. Enlarged hearts in both female (left panel) and male (right panel) *Rhbdf1*^*−/−*^ mice. Enlarged hearts were observed in pups older than three weeks of age. When we measured heart and body weights of *Rhbdf1*^*−/−*^ (n=3 females; n=4 males) and *Rhbdf1*^*+/−*^ (n=4 females; n=5 males) littermates at postnatal day 21, the heart-to-body-weight ratio (HW/BW) was significantly higher *Rhbdf1*^*−/−*^ mice relative to *Rhbdf1*^*+/−*^ littermates. Data represent mean ± S.D; *p<0.05, ***p<0.001. h. Cardiac complications in *Rhbdf1*^*−/−*^ mice on postnatal day 21. Masson’s trichrome staining of heart sections from a three-week-old *Rhbdf1*^*−/−*^ mouse showing interventricular-septum and left-ventricular fibrosis. Notably, both perivascular and interstitial fibrosis were observed in *Rhbdf1*^*−/−*^ mice.

In addition to the generalized growth deficiency of *Rhbdf1*^*−/−*^ survivors, specific abnormalities were observed in the brain and heart. Gross examination of whole brains from P21 *Rhbdf1*^*−/−*^ mice revealed hemorrhages and histological examination of *Rhbdf1*^*−/−*^ brain sections confirmed several hemorrhages throughout the brain. These were evident in the ventricles adjacent to the brain stem, hippocampus, and hypothalamus (Fig. 2f), without significant inflammation. Additionally, P21 *Rhbdf1*^*−/−*^ mice displayed left atrial enlargement with left ventricular hypertrophy, which is reflected in the higher heart weight/body weight ratio in these homozygous mice compared to heterozygotes (Fig. 2g). Histological sections of the hearts revealed perivascular and interstitial fibrosis in the left ventricle and interventricular septum (Fig. 2h). In agreement with these data, heart-to-body-weight ratios revealed significant cardiac hypertrophy in the *Rhbdf1*^*−/−*^ animals. In contrast, no lesions were detected in hearts from *Rhbdf1*^*+/−*^ mice aged up to one year. Our findings in the present study—that *Rhbdf1*^*−/−*^ mice on a pure inbred C57BL/6N strain background have significantly lower body weight compared with heterozygous-null mice, with most *Rhbdf1*^*−/−*^ mice dying postnatally from brain hemorrhages and cardiac fibrosis—are similar in many ways to those of the Christova et al. study, but very different from than those of Li et al. Notably, Christova et al. generated mice on two different strain backgrounds—C57BL/6J and 129S6/SvEvTac—and showed that whereas homozygotes on each background showed severe weight loss, brain hemorrhage, and cardiac infarction, and died by ~6 weeks, heterozygotes on each background were normal. Collectively, these data suggest an essential role for RHBDF1 during postnatal development, and hereafter the definitive-null *Rhbdf1* homozygous-null mice are referred to as *Rhbdf1*^*null/null*^ mice.

Because the *Rhbdf1* and *Rhbdf2* genes have significant homology (~80%) in their sequences, we sought to address whether the phenotype of *Rhbdf1:Rhbdf2* double KO mice is more severe than that of *Rhbdf1*^*null*^ mice; i.e., embryonic or perinatal lethality. Notably, *Rhbdf2*^*−/−*^ mice are viable and fertile, and do not display growth retardation or brain and heart defects [19–21]. Interestingly, in contrast to the viability of *Rhbdf1*^*null/null*^ embryos, the combined loss of *Rhbdf1* and *Rhbdf2* resulted in sub-viability. Of 64 live births from *Rhbdf1*^*+/null*^ *Rhbdf2*^*−/−*^ parents, there were four pups born that were homozygous-null for both genes (*Rhbdf1*^*null/null*^ *Rhbdf2*^*−/−*^) (Supplementary Fig. 2a), suggesting subviability due to deficiency of both RHBDF1 and RHBDF2, and suggesting functional redundancy between RHBDF1 and RHBDF2 during embryonic development. Moreover, histopathological analysis of *Rhbdf1*^*null/null*^ *Rhbdf2*^*−/−*^ mice revealed an ‘eyelids open at birth’ phenotype (Supplementary Figs. 2b and 2c), suggestive of deficiencies in epidermal growth factor receptor (EGFR) signaling pathways [22, 23]. These findings—that the majority of *Rhbdf1*^*null/null*^ *Rhbdf2*^*−/−*^ mice die in utero—are similar to those of the study of Christova et al., who found that *Rhbdf1:Rhbdf2* double KO mice on either the C57BL/6J or the 129S6/SvEvTac backgrounds results in embryonic lethality.

### CRISPR/Cas9-mediated deletion of the critical exon(s) in *Rhbdf1* results in viable and fertile homozygous mutants with the A7 allele

We decided to investigate whether deletion of *Rhbdf1* critical exons 2 and 3, which contain the translation initiation codon, rather than deletion of the entire protein coding sequence of the target gene (as in the A5 allele), would result in mice that mimic *Rhbdf1*^*null/null*^ mice (Fig. 2) or mice that exhibit an altered phenotype owing to creation of an N-terminally truncated abnormal protein product. We used CRISPR/Cas9 to generate this *Rhbdf1* mutant allele (A7), which excised exons 2 and 3 by the NHEJ repair pathway (Fig. 3a and Supplementary Fig. 3). Mutant mice that are homozygous for this allele are hereafter referred to as viable (*Rhbdf1*^*v/v*^) mice. We observed that whereas heterozygous-viable (*Rhbdf1*^*+/v*^) mice have one wildtype band (1,300-bp PCR product) and one mutant band (728-bp), homozygous-viable (*Rhbdf1*^*v/v*^) mice have only the mutant band (728-bp) (Fig. 3b, left). The deletion of DNA sequence was further validated by Sanger’s sequencing (Fig. 3b, right). None of the *Rhbdf1*^*v/v*^ pups showed gross deformities or malformations of major organs in contrast to the majority of *Rhbdf1*^*null/null*^ mice, which were found dead by P14 (Fig. 2). *Rhbdf1*^*v/v*^ mice were healthy, viable, and fertile (Fig. 3c, 3d, and 3e). Additionally, no specific abnormalities were observed in the brain, heart, kidney, spleen, skin, or liver by histological examination (Fig. 3f and 3g and Supplementary Fig. 4a). This is also reflected in the normal heart weight/body weight ratio in *Rhbdf1*^*+/v*^ and *Rhbdf1*^*v/v*^ mice (Fig. 3g), compared with that of *Rhbdf1*^*null/null*^ mice. These findings are similar in many ways to those of the Li et al. study, who found that the *Rhbdf1* mutant mice they generated were indistinguishable from their littermate controls[18].

**Figure 3.**
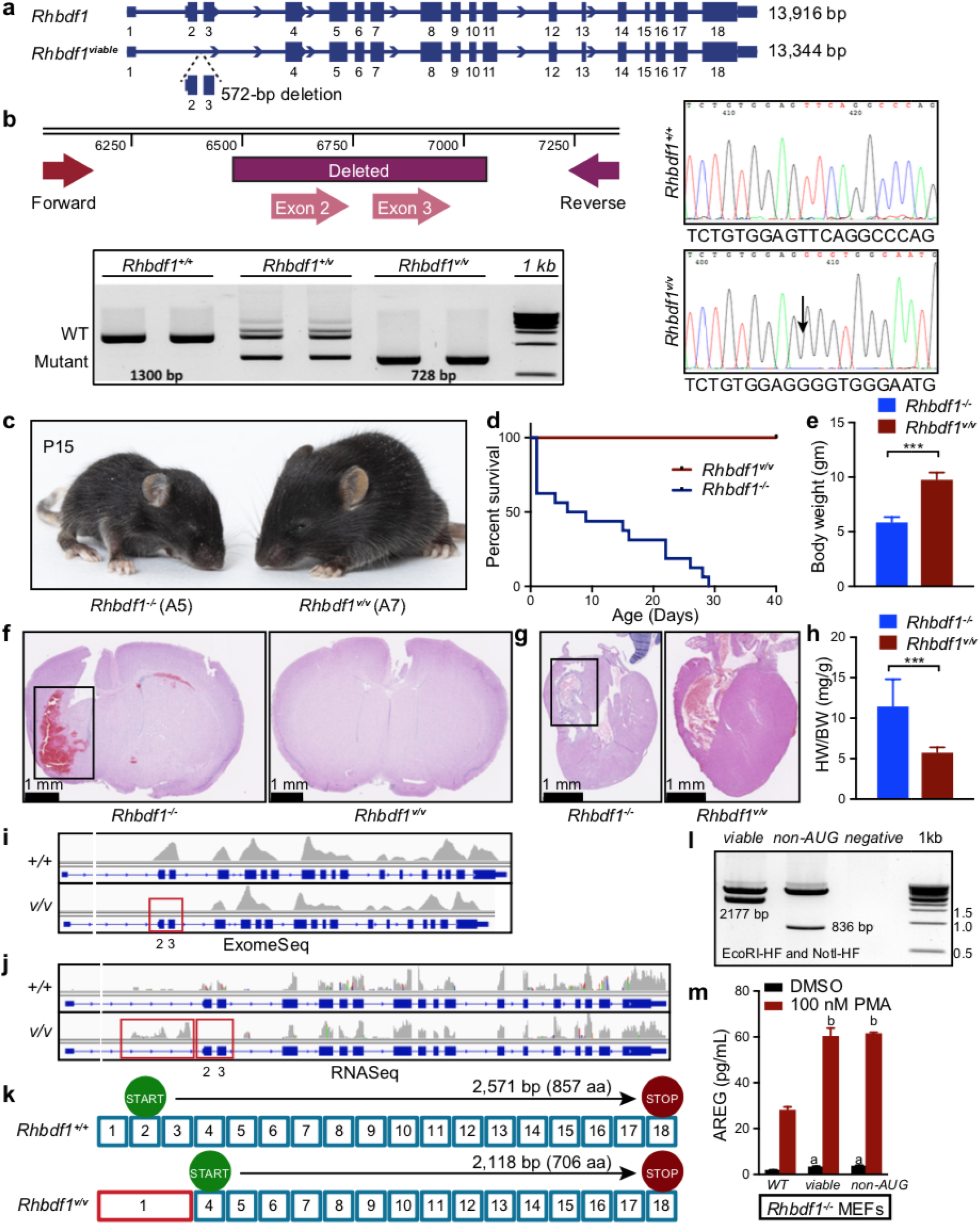
Deletion of critical exons 2 and 3 in *Rhbdf1* results in viable and healthy mice (A7) a. CRISPR/Cas9-mediated deletion of critical exons 2 and 3 resulted in an in-frame 572-bp deletion in the *Rhbdf1* gene (hereafter referred to as viable mice, *Rhbdf1*^*v/v*^). b. The schematic shows the deleted exons and the PCR strategy to amplify the region. PCR of genomic DNA with primers flanking exons 2 and 3 shows that wildtype mice have one wildtype band (1,300-bp PCR product), heterozygous mice carry a wildtype band (1,300 bp) and a mutant band (728 bp), and homozygous-null mice carry one mutant band (728 bp). Samples were run in duplicate. DNA sequencing further validated an in-frame 572-bp deletion in genomic DNA (right). Also, see Supplementary Fig. 4. c. Images of representative postnatal day 15 *Rhbdf1*^*−/−*^ mice (left) and *Rhbdf1*^*v/v*^ mice (right). d. Kaplan-Meier survival analyses showing that the lifespan of *Rhbdf1*^*v/v*^ mice (A7) is significantly longer than that of *Rhbdf1*^*−/−*^ mice (A5). e. Body weights of *Rhbdf1*^*v/v*^ mice compared with those of *Rhbdf1*^*−/−*^ mice on postnatal day 21. Data represent mean ± S.D; ***p<0.001. f. H&E-stained coronal sections of a postnatal day 21 *Rhbdf1*^*−/−*^ brain showing prominent hemorrhages in the brain; in contrast, there was no evidence of brain hemorrhaging in *Rhbdf1*^*v/v*^ mice. g. Cardiac abnormalities were observed in *Rhbdf1*^*−/−*^ mice but not in *Rhbdf1*^*v/v*^ mice (right). The black rectangle illustrates cardiac fibrosis in *Rhbdf1*^*−/−*^ mice (left). h. Also, heart weight was significantly lower in *Rhbdf1*^*v/v*^ mice than in *Rhbdf1*^*−/−*^ mice. Data represent mean± S.D; ***p<0.001. i. Exome-Seq showing loss of exons 2 and 3 in *Rhbdf1*^*v/v*^ mutant mice and not in *Rhbdf1*^*+/+*^ mice. The red rectangle shows the exons lost in the mutant mice. j. RNA sequencing analysis of spleens isolated from *Rhbdf1*^*v/v*^ mice indicating that CRISPR/Cas9-mediated deletion of critical exons 2 and 3 does not result in nonsense-mediated decay of mutant *Rhbdf1*^*v/v*^ mRNA. Note the boxed region preceding the lost exons 2 and 3 indicates the presence of alternative transcripts. k. Although the translation initiation site in exon 2 is deleted in *Rhbdf1*^*v/v*^ mice, sequence analysis of *Rhbdf1*^*v/v*^ cDNA revealed that the next in-frame translation initiation site (ATG) was in exon 4, which could potentially result in an N-terminally truncated protein to rescue the severe pathology of *Rhbdf1*^*−/−*^ mice (A5). l. Double digests of C-terminal Myc-DDK-tagged *Rhbdf1*^*viable*^ and *Rhbdf1*^*non-AUG*^ vectors using EcoRI-HF and NotI-HF restriction enzymes. m. Rescue experiments in *Rhbdf1*^*−/−*^ MEFs. RHBDF1 deficiency suppresses stimulated secretion of AREG in primary MEFs (see supplementary Figure 5); hence, we performed rescue experiments to restore stimulated AREG secretion in *Rhbdf1*^*−/−*^ MEFs (A5); *Rhbdf1*^*−/−*^ MEFs were transiently transfected with 2 μg of either a full-length vector, or a *Rhbdf1*^*viable*^ vector, or a *Rhbdf1*^*non-AUG*^ vector, using Lipofectamine LTX. 48 h post-transfection, cells were stimulated overnight with either DMSO or 100 nM PMA, and cell-culture supernatants were analyzed using a mouse AREG ELISA kit. Data represent mean ± S.D; a, significantly different from a full-length vector transfected and DMSO stimulated cells; b, significantly different from a full-length vector transfected and PMA stimulated cells.

Exome-Seq of *Rhbdf1*^*v/v*^ mice validated an in-frame 572-bp deletion in the *Rhbdf1* gene, resulting in loss of exons 2 and 3 (Fig. 3i). To examine whether this deletion produces an alternative *Rhbdf1* transcript, we performed RNA-Seq on RNA derived from spleens of *Rhbdf1*^*v/v*^ mice, which showed the *Rhbdf1*^*v/v*^ transcript lacked exons 2 and 3, but contained all of the other exons (1, and 4 through 18). Because the normal translation initiation site (ATG) in exon 3 is missing, we predicted that the *Rhbdf1*^*v/v*^ transcript could produce a protein variant or a truncated RHBDF1 protein by utilizing the next in-frame translation initiation site in exon 4. Notably, *Rhbdf1* transcript variants X2, X3, and X4 all lack exons 2 and 3 [24], but generate an alternative mRNA transcript, which is likely to result in a 706-amino acid product utilizing the next in-frame ATG in exon 4 (Fig. 3k and Supplementary Fig. 4b).

To determine whether the *Rhbdf1*^*v/v*^ truncated protein is functional and whether it could rescue RHBDF1 deficiency, we performed *in vitro* stimulated-secretion assays in primary mouse embryonic fibroblasts (MEFs). Previous studies have shown that RHBDF1 regulates protein kinase C activator phorbol ester (PMA)-mediated stimulated secretion of epidermal growth factor receptor (EGFR) ligands, including amphiregulin (AREG)[20]. Here, using PMA-induced stimulated secretion assays in *Rhbdf1*^*+/+*^ and *Rhbdf1*^*−/−*^ primary MEFs (A5 allele), we found that, upon stimulation with 100 nM PMA, *Rhbdf1*^*+/+*^ MEFs showed a significant increase in AREG secretion compared with *Rhbdf1*^*−/−*^ MEFs, indicating that RHBDF1 deficiency suppresses AREG secretion (Supplementary Fig. 5). Hence, we asked whether the variant protein can rescue the ‘stimulated secretion of AREG’ phenotype, by transiently transfecting *Rhbdf1*^*v/v*^-expressing vector in *Rhbdf1*^*−/−*^ primary MEFs, and measuring AREG levels following stimulation with PMA (Fig. 3l). Stimulation with PMA significantly increased AREG levels in *Rhbdf1*^*v/v*^-transfected MEFs compared with empty vector-transfected MEFs, suggesting that *Rhbdf1*^*v/v*^ is capable of inducing AREG secretion and thereby rescuing the phenotype.

Because a majority of double mutant *Rhbdf1*^*null/null*^ *Rhbdf2*^*−/−*^ mice die *in utero*, we asked whether the *Rhbdf1*^*v/v*^ mutation can rescue the subviability of *Rhbdf1*^*null/null*^ *Rhbdf2*^*−/−*^ mice, by generating *Rhbdf1*^*v/v*^ *Rhbdf2*^*−/−*^ mice. In contrast to the subviability of *Rhbdf1*^*null/null*^ *Rhbdf2*^*−/−*^ mice, *Rhbdf1*^*v/v*^ *Rhbdf2*^*−/−*^ mice were viable, healthy, and fertile (Supplementary Fig. 6a). Nevertheless, *Rhbdf1*^*v/v*^ *Rhbdf2*^*−/−*^ mice exhibited wavy-coated hair and ‘eyelids open at birth’ phenotypes (Supplementary Fig. 6b and 6c), indicative of alterations in epidermal growth factor receptor (EGFR) signaling pathways. Histopathological analysis revealed no evidence of heart or brain abnormalities (Supplementary Fig. 6d).

Literature suggests that, in addition to the canonical AUG start codons, non-AUG start codons are capable of initiating translation [25–28]. Notably, *Rhbdf1* protein-coding transcript 206 [24], utilizes a non-AUG start codon, CGC (arginine, R), rather than AUG (methionine, M) (Supplementary Fig. 7a). We next sought to examine whether mRNA translation initiation from non-AUG start codons can generate functional protein-coding transcripts that rescue the phenotype. Since there are no validated RHBDF1-specific antibodies, we obtained C-terminal Myc-DDK tagged expression vector for the non-AUG mutant transcript, and heterologously expressed them in 293T cells. Western blotting revealed that non-AUG vector, based on their molecular masses, generated expected size bands (~30-kDa) while transfection with an empty tagged vector did not produce any bands (Supplementary Fig. 7b). Additionally, when the non-AUG-expressing vector was transfected into *Rhbdf1*^*−/−*^ primary MEFs, it induced significant AREG secretion (Fig. 3l). This result is similar to the *Rhbdf1*^*v/v*^ transcript in transfected MEFs, suggesting that the non-AUG start codon CGC is capable of generating a functional protein-coding transcript. Thus, we conclude that mutant *Rhbdf1*^*v/v*^ protein products rescue the severe phenotype observed in mice deficient in RHBDF1 (both Christova et al. and our *Rhbdf1*^*null/null*^ mice), because of the use of alternative AUG and non-AUG start codons that nevertheless give rise to mostly functional proteins.

### Targeted KO-first allele *Rhbdf1* homozygous mutants (A3) express truncated protein isoforms that rescue the pathology of *Rhbdf1*^*null/null*^ mice

Additional crosses of mice carrying the *Rhbdf1* KO-first allele to Flp transgenic mice, followed by crosses to Cre transgenic mice, resulted in excision of the cassette flanked by the FRT sites and of exons 4-11 flanked by the *loxP* sites, respectively (Fig. 4a). These homozygous mutant mice were generated by Li et al., and are hereafter referred to as viable 2 mice (*Rhbdf1*^*v2/v2*^). Exome-Seq of these mice validated deletion of exons 4-11 in the *Rhbdf1* gene (Fig. 4b). RT-PCR (Fig. 4c and Supplementary Fig. 8) and RNA-Seq (Fig. 4d) confirmed that exons 4 through 11 are missing in the transcript, while exons 12 through 18 remain intact. The structure of the *Rhbdf1*^*v2/v2*^ transcript suggests that exons 3 and 12 might be spliced together. To test this, we performed RT-PCR on the *Rhbdf1*^*v2/v2*^ transcript using probes in exon 3 and in exon 12. Instead of an expected 209-bp product suggesting that exons 3 and 12 are joined together, we detected a 324-bp PCR product, indicating insertion of a short 115-bp DNA fragment. Sanger sequencing of this product revealed that the *Rhbdf1*^*v2/v2*^ mRNA retained the *En2* splice acceptor sequence (Supplementary Figs. 8b,c) from the cassette (Supplementary Fig. 8e). Nonetheless, the resultant *Rhbdf1*^*v2/v2*^ mutant transcript retains an in-frame termination codon (TAA), which will result in a 10.8-kDa short truncated protein product (Supplementary Fig. 8f).

**Figure 4.**
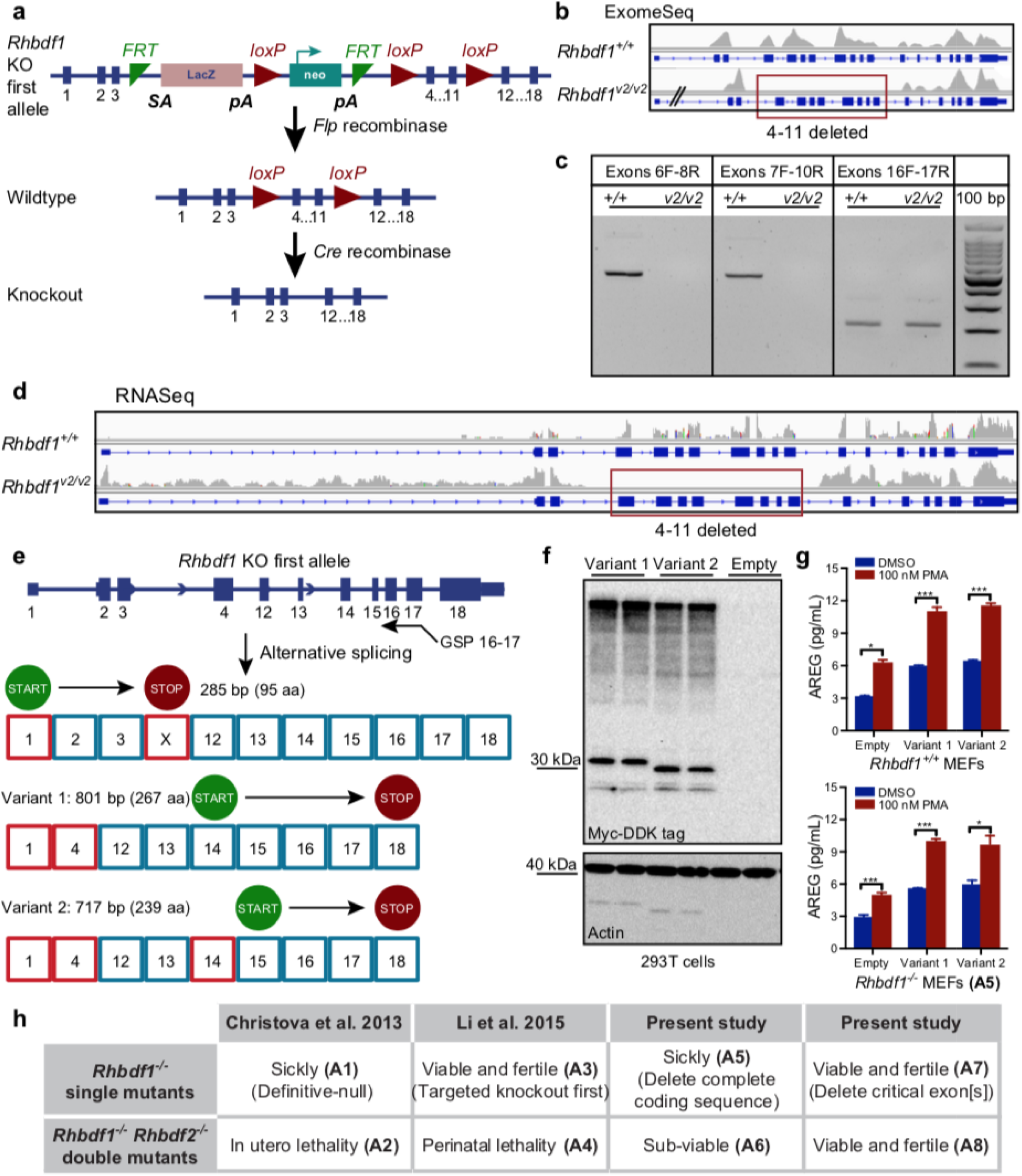
Targeted KO-first targeting strategy in *Rhbdf1* (A3) generates novel transcripts and N-terminally truncated functional proteins. a. Schematic of the strategy used by Li et al. for generation of *Rhbdf1*^*−/−*^ homozygous mutant mice; the *Rhbdf1* KO-first allele was crossed to Flp recombinase mice to remove the FRT-flanked “lacZ reporter and a neomycin resistance (neo) gene” to generate conditional-ready mice, which were later crossed with cre transgenic mice to excise the floxed gene segment (exons 4-11), generating *Rhbdf1*^*−/−*^ homozygous mutant mice (hereafter referred as viable2 mice, *Rhbdf1*^*v2/v2*^ mice). b. Whole-exome sequencing of spleen tissue from *Rhbdf1*^*v2/v2*^ mice showing loss of exons 4 through 11 in *Rhbdf1*^*v2/v2*^ mutant mice. c. RT-PCR on spleens from *Rhbdf1*^*+/+*^ and *Rhbdf1*^*v2/v2*^ mutant mice using primers to amplify exons 6 through 8, exons 7 through 10, and exons 16 and 17. Exons 4-11 are deleted in *Rhbdf1*^*v2/v2*^ mutant mice; hence no amplicons were generated using either exon 6 forward and exon 8 reverse, or exon 7 forward and exon 10 reverse, primers. However, exon 16 forward and exon 17 reverse primers generated a 211-bp product. d. RNA-Seq analysis of spleens from *Rhbdf1*^*v2/v2*^ mutant mice indicating loss of exons 4 through 11; however, there is strong evidence for mutant mRNA, as indicated by the presence of the rest of the transcript, which encodes exons 12 through 18 and is not degraded by the nonsense-mediated decay mechanism. e. Schematic representation of exons and introns in the *Rhbdf1*^*v2/v2*^ mutant allele. 5’ RACE using a gene-specific exon 16-17 fusion primer (GSP) was used to obtain 5’ ends of the *Rhbdf1*^*v2/v2*^ mutant mRNA. We identified several novel mutant mRNAs with different translation initiation sites that could potentially generate N-terminally truncated RHBDF1 mutant proteins. See supplemental figures for variant protein and 5’ UTR sequences. Alternative exons are indicated as red boxes; predicted translation initiation sites are indicated by “START,” and termination codons are indicated by “STOP.” f. C-terminal Myc-DDK-tagged *Rhbdf1*^*v2/v2*^ variant protein 1 (lanes 1, 2) or variant protein 2 (lanes 3,4), or empty vector (lanes 5, 6) were transiently expressed in 293T cells, and cell lysates were analyzed using western blotting with FLAG-specific antibody. After visualization of blots with a G:Box chemiluminescent imaging system, blots were washed, blocked in 5% nonfat dry milk, and re-probed with anti-actin antibody. g. Rescue of phenotype in *Rhbdf1*^*−/−*^ MEFs. *Rhbdf1*^*+/+*^ (top) and *Rhbdf1*^*−/−*^ (bottom) MEFs were transiently transfected with 2 μg of either variant 1 or variant 2 vectors, or an empty vector, using Lipofectamine LTX. 48 h post-transfection, cells were stimulated overnight with either DMSO or 100 nM PMA, and cell-culture supernatants were analyzed using a mouse AREG ELISA kit. Data represent mean ± S.D; *p<0.05, ***p<0.001. h. Comparison of phenotypes of previously generated *Rhbdf1* single mutant and *Rhbdf1:Rhbdf2* double mutant mice with those of mice generated in the present study. The viability of *Rhbdf1*^*v/v*^, *Rhbdf1*^*v2/v2*^, and *Rhbdf1*^*v/v*^ *Rhbdf2*^*−/−*^ double mutant mice demonstrates that the incomplete coding sequences not degraded by NMD have functional significance, and, moreover, that targeted KO-first conditional mutagenesis (A3) or CRISPR/Cas9-mediated deletion of critical exon(s) (A7), in contrast to complete deletion of exons (A1 and A5), can yield gain-of-function alleles.

We next sought to explore whether the conditional *Rhbdf1*^*v2/v2*^ mutation leads to additional truncated proteins via the use of an alternative promoter usage or via exon skipping to reinitiate translation[15, 29, 30]. To evaluate the 5’ ends of mutant *Rhbdf1*^*v2/v2*^ mRNA transcripts, we performed 5’ RACE on first-strand cDNA using a gene-specific exon 16-17 fusion primer, based on the RT-PCR evidence for the presence of a 211-bp product (Fig. 4c). The resulting RACE products were characterized by Sanger sequencing following agarose gel electrophoresis and In-Fusion cloning. Sequence analyses of identified clones revealed the existence of an alternative exon 1, 199 bp upstream of the conventional exon 1 (Supplementary Figs. 9,10). Interestingly, we found evidence for two potential transcripts that utilize this alternative exon 1 (Supplementary Figs. 11, 12) that could generate N-terminally truncated *Rhbdf1*^*v2/v2*^ proteins by alternative splicing (Fig. 4e). We also obtained C-terminal Myc-DDK-tagged expression vectors for variant 1 and 2 mutant transcripts, and heterologously expressed them in 293T cells (Supplementary Fig. 13). Western blotting revealed that both variants (1 and 2), based on their molecular masses, generated expected size bands (~32-kDa and ~29-kDa) while transfection with an empty tagged vector did not produce any bands (Fig. 4f).

Lastly, using PMA-induced stimulated secretion assays, we observed that, first, in both *Rhbdf1*^*+/+*^ and *Rhbdf1*^*−/−*^ MEFs, transfection alone with either variant protein 1 or 2 significantly increased AREG secretion compared with that of the empty vector (Fig. 4g). Second, stimulation with PMA further increased AREG levels in both variant protein 1- or 2-transfected MEFs compared with empty vector transfected MEFs, suggesting that both of the N-terminally truncated RHBDF1 variant proteins are capable of inducing AREG secretion and thereby rescuing the phenotype (Fig. 4g). Collectively, these data suggest that the lack of spontaneous pathological phenotypes in the *Rhbdf1*^*v2/v2*^ mutant mice could be explained by the presence of residual mRNA transcripts and protein isoforms that can prevent the more severe phenotype observed in the *Rhbdf1*^*null/null*^ mice, and that the *Rhbdf1*^*v2/v2*^ mice with the targeted KO-first allele (A3) are not homozygous-null.

### Identifying unexpected translation by analyzing incomplete protein-coding sequences (CDS) evading nonsense-mediated decay

We next investigated whether we could find mouse and human protein-coding genes that are likely predisposed to targeted KO-first- and CRISPR/Cas9-mediated illegitimate or unexpected transcription and translation. Several factors, including exon skipping, alternative promoter usage, and translation reinitiation at AUG and at non-AUG start codons, could result in abnormal phenotypes. Identifying mouse and human protein-coding genes that express incomplete coding transcripts (Supplementary Tables 1-4) that are not subjected to NMD could help identify genes resilient to targeted KO-first or frameshift indels. We found that at least 35% of mouse and 45% of human protein-coding genes generate protein-coding transcripts that are not eliminated by NMD (Fig. 5a). For instance, the Ensembl site lists nine transcripts (*Rhbdf1-*201 through - 209) for mouse *Rhbdf1* (Fig. 5b). *O*f the four protein-coding *Rhbdf1* transcripts (201, 206, 207, and 209), transcript 206, which has a 5’ incomplete CDS and is comprised of only exons 14 through 18. As such, this transcript is not likely to be degraded by NMD when alleles with targeting event 5” of these exons are generated. Indeed, our tests in the present study were focused on amplifying exons 15 through 18 in the *Rhdbf1*^*v2/v2*^ allele (Fig. 4c). Additionally, 5’ RACE on first-strand *Rhdbf1*^*v/v*^ and *Rhdbf1*^*v2/v2*^ cDNA was performed using a gene-specific exon 16-17 fusion primer (Fig. 4e). Importantly, targeted KO-first mutagenesis excised only exons 4 through 11, leaving the exons in the *Rhbdf1*-206 transcript (14 to 18) untouched and thus not likely to be subject to NMD.

**Figure 5.**
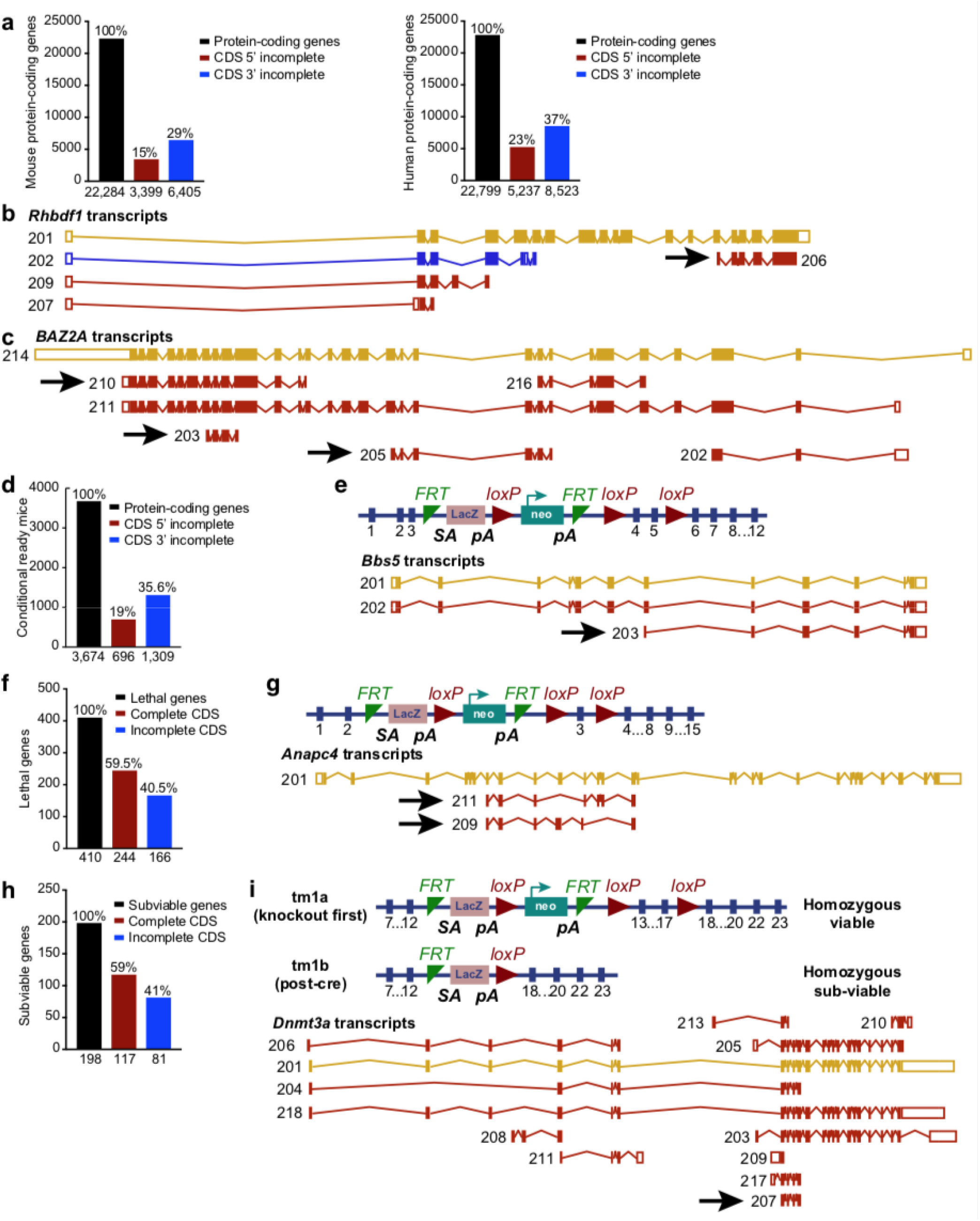
An ‘incomplete protein-coding CDS’ approach to identifying on-target unintended transcription and translation reinitiation. a. Bioinformatics analyses of mouse (left) and human (right) protein-coding genes with incomplete 5’ CDSs and incomplete 3’ CDSs reveals that ~50% of genes are potentially capable of generating shorter transcripts and truncated proteins. The physiological significance of these incomplete CDSs is unclear. However, it is yet to be determined whether gene targeting strategies must be designed to ensure the deletion of these incomplete, yet existing coding sequences that are not degraded by NMD. b. The various transcripts of the mouse *Rhbdf1* gene listed in Ensembl. In this scenario, the definitive null design, rather than either the targeted KO-first or CRISPR/Cas9-mediated deletion of critical exon(s), would prevent expression of a short 5’ incomplete *Rhbdf1*-206 transcript. c. CRISPR/Cas9-mediated frameshift mutation failed to induce complete deficiency of *BAZ2A* in HAP1 cells, however, based on the presence of 5’ and 3’ incomplete CDSs (arrows), *BAZ2A* is likely to generate truncated proteins upon targeted KO-first- or CRISPR/Cas9-induced on-target mutagenesis. d. Bioinformatics analyses of 3,674 conditional-ready mouse strains identified that nearly 55% of conditional ready genes generate incomplete 5’ and 3’ CDSs. e. Schematic of *Bbs5* KO-first allele in which the ‘lacZ reporter gene and neomycin resistance gene’ flanking FRT recognition sites is placed downstream of the third exon of the gene, and critical exons 4 and 5 are floxed by *loxP* sites. Following cre-mediated recombination, despite missing exons 4 and 5, *Bbs5* 5’ incomplete CDS 203 may generate an N-terminally truncated protein. f. Bioinformatics analyses of 410 lethal IMPC mouse strains identified that 166 essential genes that generate incomplete 5’ and 3’ CDSs. g. Schematic of *Anapc4* KO-first allele in which the ‘lacZ reporter gene and neomycin resistance gene’ flanking the FRT recognition sites is placed downstream of the second exon of the gene, and the critical exon 3 is floxed by *loxP* sites. Although cre recombination deletes critical exon 3, translation re-initiation from incomplete CDSs 209 and 211 may rescue the lethal phenotype of *Anapc4* homozygous-null mice. h. Analyses of 198 subviable IMPC mouse strains identified 81 genes generating incomplete 5’ and 3’ CDSs. i. The targeted KO-first strategy in *Dnmt3a* results in conflicting phenotypes. Whereas the *tm1a* KO-first allele generates homozygous viable mice, the *tm1b* allele generates homozygous subviable mice (top). Multiple protein-coding transcripts of *Dnmt3a* are shown. The *Dnmt3a* 207 protein-coding transcript has incomplete 5’ and 3’ CDSs (arrow) (bottom).

*BAZ2A* is another example of a gene that was recently identified during systematic characterization of frameshift KO mutations in HAP1 cells showing residual protein expression [15], that we could identify here based on the presence of incomplete CDS (Fig. 5c). In Supplementary Table 5, we present additional examples of the genes that retain residual protein expression/function upon CRISPR/Cas9-induced indels in recent literature [14–17, 28–31] and that we validate as good candidates for unanticipated translation based on the incomplete CDS approach.

We also analyzed 3,674 conditional-ready mouse strains [32] using the incomplete CDS approach (Supplementary Table 6), and determined that at least 40% of conditional-ready alleles can skew the mouse phenotype (Fig. 5d). For instance, the *Bbs5* conditional-ready allele is likely to generate a 5’ incomplete CDS and an N-terminally truncated protein despite missing exons 4 and 5, which are lost upon cre-mediated recombination (Fig. 5e).

In contrast to mutation of gene function via unsolicited translation following CRISPR/Cas9 or targeted KO-first gene targeting, can unsolicited translation be advantageous for exploring the function of genes that are essential for survival? The International Mouse Phenotyping Consortium (IMPC) systematically characterized 5,000 KO mouse strains and identified 410 lethal and 198 subviable genes [33]. Here, we analyzed all of the lethal and subviable genes (Supplementary Table 6) and found that, upon cre-recombination of conditional-ready alleles, nearly 40% of the lethal genes and 40% of the subviable genes may result in unwanted translation and could potentially partially rescue the mouse phenotype generated by a null mutation (Fig. 5f and 5g). We show an example of the *Anapc4* KO-first allele, in which the “lacZ reporter gene and neomycin resistance gene” flanking the FRT recognition sites for FLP recombinase is placed downstream of the second exon of the gene, and the critical exon 3 is floxed by *loxP* sites for recognition by Cre recombinase. Flp recombinase removes the lacZ and neomycin cassette, allowing reversal to a conditional-ready allele, while Cre recombinase deletes the critical exon, presumably generating null or conditional mutants (Fig. 5h). However, *Anapc4* expresses incomplete protein-coding transcripts that lack the critical exon 3 and that evade NMD (*Anapc4*-209 and −211), suggesting that these protein-coding transcripts could rescue the phenotype and impact interpretation of the phenotype.

In accordance with our observations, emerging evidence from IMPC suggests that several mouse strains generated using the targeted KO-first strategy exhibit conflicting phenotypes: *Dnmt3a* (*tm1a* allele is homozygous viable; *tm1b* allele is homozygous subviable) (Fig. 5i); *Foxj3* (*tm1a* is homozygous viable; *tm1b* is homozygous lethal); and *Slc20a2* (*tm1a* is homozygous viable; *tm1b* is homozygous subviable), suggesting at least partial rescue of gene function due to residual protein activity (Table 1 and Supplementary Table 7). Notably, both the *tm1a* and *tm1b* mouse strains, KO-first and post-cre, respectively, should exhibit the same phenotype. Thus, we strongly believe that, despite rigorous quality control of mutant mouse strains by the IMPC, additional post-genomic-DNA characterization is required to ensure elimination of any residual proteins of the targeted allele that might interfere with the phenotype. This is particularly important when novel sequences are incorporated, such as those introduced by targeted KO-first, making it extremely difficult to predict any novel mRNA transcripts that may be generated.

**Table 1.** The KO-first gene-disruption strategy results in conflicting phenotypes in homozygous mutant mice

| Mice generated using<br>KO-first strategy | Phenotype |  |  |
| --- | --- | --- | --- |
|  | Homozygous<br>viable | Homozygous<br>subviable | Homozygous<br>lethal |
| *Bbox1* * | *em1* | *tm1b* |  |
| *Caskin1* | *em1* | *tm1b* |  |
| *Cpsf3* * |  | *tm1b* | *tm1a* |
| *Dbn1* * | *em1* | *tm1b* | *tm1b* |
| *Dnmt3a* * | *tm1a* | *tm1b* |  |
| *Dppa1* |  | *tm1b* |  |
| *Exoc3l2* | *tm1a* |  | *tm1b* |
| *Foxj3* * | *tm1a* |  | *tm1b* |
| *Ino80* |  | *tm1b* | *tm1a* |
| *Itm2c* * | *em2* | *tm1b* |  |
| *Kdm8* * |  | *tm1a* | *tm1b* |
| *Lmna* * |  | *tm1a* | *tm1b* |
| *Nxn* * |  | *tm1a* | *tm1b* |
| *Rundc1* * |  | *tm1b* | *tm1a* |
| *Slc20a2* * | *tm1a* | *tm1b* |  |
| *Tox3* * | *em1* | *tm1b* |  |
| *Zranb1* | *tm1a* | *tm1b* |  |

Analysis of 427 conditional-ready mouse strains generated using the KO-first strategy reveals that at least 4% of mouse strains show conflicting phenotypes. \*Notably, 12 of 17 lethal/subviable genes express incomplete protein-coding sequences that could generate truncated proteins to potentially skew the phenotype of homozygous mutant mice.

## Discussion

Several recent studies have reported that CRISPR/Cas9-mediated on-target gene KO can produce novel unintended mRNA isoforms and truncated proteins that could at least partially rescue activity of the targeted gene; Lalonde and colleagues found that despite CRISPR/Cas9-induced on-target frameshift mutations in the phosphatase and actin regulator 1 *(PHACTR1)* gene in human aortic endothelial cells, exon skipping results in novel functional *PHACTR1* transcripts [29]. Also, in zebrafish lines, CRISPR/Cas9-generated KOs display irregular phenotypes, owing to activation of cryptic splice sites in mutant mRNA transcripts [17]. Similarly, to examine whether CRISPR/Cas9 mutations induce exon skipping to generate in-frame transcripts and truncated proteins with functional outcomes, Mou et al. targeted exon 3 of the beta-catenin gene *Ctnnb1* in a cell line, which induced indels as expected. However, unexpected splicing together of exons 2 and 4, or of exons 2 and 5, together resulted in novel protein isoforms [30]. Immunoblotting with beta-catenin specific antibody revealed the presence of mutant proteins lacking exon 3-encoded amino acid residues, suggesting that indels induced by CRISPR/Cas9-sgRNAs can lead to skipping of edited exons, forcing production of truncated proteins. Makino et al. also found that biallelic on-target frameshift mutations in the zinc finger protein-encoding gene *Gli3* did not result in homozygous null alleles and owing to illegitimate translation, GLI3 protein expression persisted in mouse NIH3T3 cells[14]. More recently, Kapahnke et al. reported that editing of the flotillin-1 gene (*FLOT1*) in HeLa cells resulted in translation of an aberrant protein with a dominant negative effect due to alternative splicing [31]. A more comprehensive analysis of residual protein expression in HAP1 cells following CRISPR/Cas9-induced frameshift mutations by Smits and colleagues further supports the observation that truncated proteins can be generated by frameshift mutations either by skipping of the edited exon or translation reinitiation [15]. Nevertheless, there have been no reports demonstrating the physiological significance of truncated protein isoforms generated by mutant mRNA transcripts to date.

Given the possibility that conflicting data with respect to phenotype can result from differences in the outcomes of different gene targeting methods [1, 18], we comprehensively characterized *Rhbdf1* mutant alleles generated by distinct gene-targeting strategies—definitive-null, CRISPR/Cas9, and targeted KO-first. Initiating with Exome-Seq, our workflow involved RT-PCR, RNA-Seq, 5’ RACE-PCR, heterologous expression of tagged expression vectors in 293T cells, and functional rescue assays. We find that whereas complete loss of exons in *Rhbdf1*^*null/null*^ mice causes growth retardation, brain and heart defects, and death of the majority of mice by P14, CRISPR/Cas9 *Rhbdf1*^*v/v*^ mice, lacking the start codon, and targeted KO-first *Rhbdf1*^*v2/v2*^ mice, lacking exons 4 through 11, do not display growth retardation or any other pathologies observed in *Rhbdf1*^*null/null*^ mice. Additionally, based on our previous findings showing that N-terminally truncated RHBDF2 causes a gain-of-function phenotype [20], and our data here showing that the *Rhbdf1*^*v/v*^ allele rescues the subviability of *Rhbdf1*^*null/null*^ *Rhbdf2*^*−/−*^ mice, causing a wavy hair coat phenotype and inducing enhanced stimulated-secretion of AREG in both *Rhbdf1*^*+/+*^ and *Rhbdf1*^*−/−*^ MEFs, we suggest that *Rhbdf1*^*v/v*^ and *Rhbdf1*^*v2/v2*^ mutant mice generate residual mRNA transcripts and protein isoforms and are gain-of-function mutations. These data also indicate that targeted KO-first alleles can generate gain-of-function alleles that rescue the severe phenotype caused by the null alleles (Supplementary Fig. 14).

Moreover, our observation that genes can skip exons with stop codons to reinitiate transcription *in vivo* to potentially rescue biological activity is in accordance with previous *in vitro* studies [14, 15, 29, 30], suggesting the need for deeper analyses of genetically modified alleles. We propose that in addition to alternative mRNA splicing, other possible mechanisms that must be accounted for while characterizing mutant mouse strains include alternative promoter usage and translation initiation from non-AUG start codons [28, 34–36], both of which can often be assessed by 5′ RACE-PCR, residual protein analysis by immunoblotting, and functional rescue assays.

Systematic generation and phenotyping of thousands of KO mouse strains by the IMPC is one of the most significant biological resources available to researchers for understanding mammalian gene function [37]. Despite advances in CRISPR-Cas genome editing technologies, generating conditional KOs is rather challenging [38], and, furthermore, targeted KO-first alleles offer a unique advantage by facilitating monitoring of gene function in a tissue- and temporal-specific manner. Nevertheless, the present study emphasizes that nature can circumvent gene targeting strategies to affect the phenotype of homozygous mutant mice. However, careful selection of gene-targeted ES clones and subsequent residual mRNA and protein analysis can considerably improve the likelihood of generating mice harboring conditional or loss-of-function alleles with pertinent phenotypes. Moreover, our data demonstrating that exon skipping in *Rhbdf1*^*v/v*^ mice prevents the multi-organ pathology observed in RHBDF1-deficient mice, and rescues embryonic lethality in RHBDF1:RHBDF2 double deficiency, indicates that the limitation of targeted KO-first and CRISPR/Cas9 approaches may provide opportunities to generate gain-of-function alleles that can either provide a partial function of a lethal gene to enable further examination of its phenotype in detail, or that can potentially be harnessed for therapeutic benefits in humans.

## Materials and Methods

### Mice

B6N(Cg)-*Rhbdf1*^*tm1.1(KOMP)Vlcg*^/J mice, stock number 026424, were obtained from The Jackson Laboratory (JAX) Repository and were bred and maintained in a barrier facility at JAX with a 12-hour light to 12-hour dark cycle. Intercrossing heterozygotes generated *Rhbdf1*^*−/−*^ mice. For generating the viable mice (*Rhbdf1*^*v/v*^), fertilized oocytes from C57BL/6J mice (stock #664) were microinjected using two sgRNAs targeting intron 1-2 and intron 3-4, respectively. Each sgRNA was delivered at a concentration of 50 ng/μl, along with Cas9 mRNA (Trilink) at 100 ng/μl and Cas9 protein (PNABio) at 30 ng/μl. The microinjection reagents were combined in nuclease-free T10E0.1 pH 7.5 (Integrated DNA Technologies; Tris 10mM, EDTA 0.1mM) supplemented with Rnasin (Promega) at 0.2 U/ul. The sgRNAs were both designed as TRU-guides with 18 nt of target-specific sequence and utilized the backbone modifications described by Chen et al[40]. The two guide target sites are shown here with PAM in parentheses: 5’-GGAGGATCTGTGGAGTTC(agg) and 5’-GAATCCCTGTTAGGTACC(agg). A total of 128 embryos were microinjected and transferred into 6 pseudopregnant females. One transfer failed to deliver any liveborn pups, but 31 mice (16F/13M) were born from the 107 embryos that were successfully transferred (29% survival). Nine of 31 (29%) pups showed a dropout allele (large deletion of the intervening sequence between the two cut sites). Genomic DNA from tails was amplified using PCR and then sequenced confirmed by Sanger sequencing. To validate germline transmission, mice harboring a 569-bp dropout were backcrossed to C57BL/6J mice and the resulting N1 offspring were intercrossed to generate the *Rhbdf1*^*v/v*^ homozygous mutant mice. Intercrossing *Rhbdf1*^*+/−*^ with *Rhbdf2*^*−/−*^ mice generated *Rhbdf1*^*−/ −*^ *Rhbdf2*^*−/−*^ double homozygous-null pups, and intercrossing *Rhbdf1*^*v/v*^ with *Rhbdf2*^*−/−*^ mice generated *Rhbdf1*^*v/v*^*Rhbdf2*^*−/−*^ double homozygous-null mice. Mice were provided food pellets (Purina LabDiet 5K52; 6% fat, autoclaved) and acidified water *ad libitum.* The Animal Care and Use Committee at JAX approved all experimental procedures.

### Histology

Mice euthanized by CO_2_ asphyxiation were gently perfused through intracardiac injection of PBS followed by injection of 4% paraformaldehyde. Tissues were fixed overnight by immersion in 10% neutral buffered formalin and then transferred to 70% ethanol. Tissue sections were stained with either hematoxylin and eosin (H&E) or Masson’s trichrome stain.

### 5’ RACE

The SMARTer RACE 5’/3’ Kit was used for performing 5’-rapid amplification of cDNA ends (RACE) using total RNA according to the manufacturer’s instructions. The integrity of total RNA isolated from spleen tissues was assessed using the Eppendorf BioSpectrometer. Total RNA (1 μg) was converted into RACE-Ready first-strand cDNA and was diluted with 10 μls of Tricine-EDTA. A 28-bp gene-specific primer (see Supplementary Figure 10) was used in touchdown PCR with 2.5 μls of 5’-RACE-Ready cDNA as template to generate the full-length cDNA. RACE products were analyzed following agarose gel electrophoresis, gel extraction, In-Fusion cloning, and Sanger sequencing.

### Cell culture

Primary mouse embryonic fibroblasts (MEFs) were isolated from embryos at 13.5 dpc (days post coitus) as previously described[20] using Liberase DL (Sigma-Aldrich). Cells were grown in a humidified chamber at 37°C with 5% CO2. MEFs and 293T cells were grown in DMEM 10% fetal bovine serum and antibiotic/antimycotic (ThermoFisher Scientific). MEFs were plated in 6-well dishes for transient transfection, western blotting, and ELISA.

### Site-directed mutagenesis and DNA cloning

Mouse expression plasmids containing cDNA encoding *Rhbdf1* was obtained from OriGene. The Q5^®^ Site-Directed Mutagenesis Kit was used to introduce site-specific mutations in the mouse *Rhbdf1* plasmid according to the manufacturer’s instructions and confirmed by Sanger sequencing. Primers were designed using the NEBaseChanger tool for site-directed substitutions and deletions, and purchased from IDT.

### Genotyping

2-mm tail tips were incubated overnight in lysis buffer containing Proteinase K at 55°C in a shaker incubator and denatured at 95°C for 5 min. 1 μl of undiluted DNA was used for 25 μls PCR reaction. For genotyping primers and protocols, see Supplementary Figure 15.

### Reverse transcriptase PCR (RT-PCR)

The Maxima First Strand cDNA Synthesis Kit (ThermoFisher Scientific) was used for cDNA synthesis from total RNA (1μg) using oligo(dT)_18_ primers. Reaction tubes containing either no reverse transcriptase or no template served as negative controls. cDNA synthesis was performed according to the manufacturer’s instructions.

### SYBR Green Real-Time PCR

PowerUp SYBR Green Master Mix was used for running the real-time PCR (ThermoFisher Scientific). For a 10 μL/well PCR reaction, 100 ng of cDNA, 800 nM exon 16 forward primer (CCTCCTGCCTTTCCTCAATC) and 800 nM exon 17 reverse primer (GAAGATGGCACTGGCTAGATT), and 5 μL of 2X master mix were mixed together and PCR was run on a ViiA 7 Real-Time PCR System using the standard cycling mode (primer T_m_≥ 60°C).

### Immunoblotting

293T cells seeded in six-well dishes and transiently transfected with expression plasmids were treated with 300 μls of RIPA lysis buffer (Cell Signaling Technology) and incubated on ice for 30 min. Cells were then centrifuged at 13,000 x g at 4°C for 10 min, and 20 μg of total cell lysate was loaded in 4-20% Tris-Glycine Mini Gels and samples were allowed to migrate for 90 min. Proteins were transferred onto a PVDF membrane using the iBlot™ Gel Transfer device, and membranes blocked with 5% milk for 1 hr at room temperature, and exposed to either FLAG-specific (Origene, #TA50011) or actin (Cell Signaling Technology, #4970) antibodies overnight at 4°C. Membranes were then washed and exposed to anti-mouse and anti-rabbit secondary antibodies, respectively, before visualizing the blots using a G:BOX gel documentation system (Syngene, Frederick, USA).

#### DNA isolation and exome sequencing

DNA was isolated from spleen using the Wizard Genomic DNA Purification Kit (Promega) according to the manufacturer’s protocols. DNA quality was assessed using Genomic DNA Screen Tape (Agilent Technologies) and Nanodrop 2000 spectrophotometer (Thermo Scientific). DNA concentration was assessed using a Qubit dsDNA BR Assay Kit (Thermo Scientific). The library was prepared by the Genome Technologies core facility at JAX using Kapa Hyper Prep (Roche Sequencing and Life Science) and SureSelectXT Mouse All Exon V2 Target Enrichment System (Agilent Technologies), according to the manufacturer’s instructions. Briefly, the protocol entails shearing the DNA using the Covaris E220 Focused-ultrasonicator (Covaris), ligating Illumina- specific adapters, and PCR amplification. The amplified DNA library is then hybridized to Mouse All Exon probes, amplified using indexed primers, and checked for quality and concentration using High Sensitivity D5000 Screen Tape (Agilent Technologies) and quantitative PCR (KAPA Biosystems), according to the manufacturers’ instructions. The library was sequenced 100-bp paired-end on the HiSeq 4000 (Illumina) using HiSeq 3000/4000 SBS Kit reagents (Illumina). Sequenced reads were filtered and trimmed for quality scores > 30 using a custom python script. Filtered Exome-Seq reads were aligned to *Mus musculus* GRCm38 using BWA-mem (0.7.9a-r786). Bam files were processed as needed (e.g., sorting, deduplication) with Picard/GATK (3.4-0).

#### RNA isolation and sequencing

RNA was isolated from spleen using the MagMAX mirVana Total RNA Isolation Kit (ThermoFisher) and the KingFisher Flex purification system (ThermoFisher). The tissue was lysed and homogenized in TRIzol Reagent (ThermoFisher). After the addition of chloroform, the RNA-containing aqueous layer was removed for RNA isolation according to the manufacturer’s protocol, beginning with the RNA bead binding step. RNA concentration and quality were assessed using the Nanodrop 2000 spectrophotometer (Thermo Scientific) and the RNA Total RNA Nano assay (Agilent Technologies). The library was prepared by the Genome Technologies core facility at JAX using the KAPA mRNA HyperPrep Kit (KAPA Biosystems), according to the manufacturer’s instructions. Briefly, the protocol entails isolation of polyA containing mRNA using oligo-dT magnetic beads, RNA fragmentation, first and second strand cDNA synthesis, ligation of Illumina-specific adapters containing a unique barcode sequence for each library, and PCR amplification. The library was checked for quality and concentration using the D5000 Screen Tape (Agilent Technologies) and quantitative PCR (KAPA Biosystems), according to the manufacturers’ instructions. The library was sequenced 100 bp paired-end on the HiSeq 4000 (Illumina) using HiSeq 3000/4000 SBS Kit reagents (Illumina). Filtered RNA-Seq reads were aligned to *Mus musculus* GRCm38 using Bowtie2 (v2.2.0).

### AREG ELISA

MEFs cultured in collagen-coated six-well plates were transiently transfected with indicated expression vectors and stimulated overnight with100 nM PMA (R&D Systems, Minneapolis, USA). Cell culture supernatant was centrifuged at 2500xg at 4°C for 5 min and 100 uls of supernatant was used to measure AREG protein levels by a Mouse Amphiregulin DuoSet ELISA Developmental Kit (#DY989, R&DSystems).

#### Incomplete human and mouse protein-coding sequences

Data were accessed from Ensembl via the SQL databases (useastdb.ensembl.org; user anonymous)[24]. The databases homo_sapiens_core_99_38 and mus_musculus_core_99_38 were queried for all transcripts where the attribute type was cds_start_NF(CDS start not found) or cds_end_NF (CDS end not found). Transcript IDs were used to select for gene ids, Ensembl stable gene ids, and biotypes assigned to transcripts and genes. Ensembl gene ids were used to select gene symbols via Ensembl Biomart. Data were collected in Microsoft Excel, and functions of Excel were used to combine (VLOOKUP) and filter the datatypes. We have developed a web application (https://incompletecds.jax.org) to identify protein-coding genes that could be predisposed to targeted KO-first- and CRISPR/Cas9-mediated unexpected transcription and translation.

### Statistical analysis

Prism v7 software (GraphPad) was used to generate the Kaplan–Meier survival curves. Student’s t-test was used to determine whether there was a statistically significant difference between two groups. P<0.05 was considered to be statistically significant.

## Authors’ contribution

VH, LDS, MVW designed and supervised the project. VH, BEL, and DL performed the experiments. GAC and VK contributed to bioinformatic analyses. VH, LDS, MVW analyzed the data and wrote the manuscript. All authors critically reviewed and approved the manuscript.

## Acknowledgements

We gratefully acknowledge the contributions of the JAX core for expert assistance with the work described in this publication, especially Mary Barter, Chrystal Snow, Christina Petros, and Heidi Munger at the Genome Technologies, Peter Kutny and Bill Buaas at Genetic Engineering Technologies, Gregg TeHennepe at Computational Sciences, and Nick Gott and Elaine Bechtel at Histopathology and Microscopy Sciences. We thank Dr. Carl P. Blobel for providing frozen spleens from the *Rhbdf1*^*v2/v2*^ mice and for critically reading the manuscript. We thank the members of the Rosenthal laboratory at JAX for technical support and for helpful suggestions. We thank Michelle L. Farley, Cindy Avery, and Todd Nason for maintenance of mouse colonies and for monitoring of early pup survival, and the ‘eyelids open at birth’ phenotype. We thank Lihong Zhao, Rosalinda A. Doty, Richard S. Maser, and Ryuta Ishimura for helpful discussion, and Stephen B. Sampson for critical reading of the manuscript. We also thank Zoe Reifsnyder in preparing illustrations for the manuscript.

**Additional files**

**Supplementary figures 1-15**

**Supplementary tables 1-7**

