## Supplemental Table 5 for "Genes adapt to outsmart gene targeting strategies in mutant mouse strains by skipping exons to reinitiate transcription and translation"

| Species | Gene(s) | 5' Incomplete | 3' Incomplete | Approach | Reference |
|---|---|---|---|---|---|
| Human | *FLOT1* | 5’ | 3' | CRISPR/Cas9 | [1] |
| Mouse | *Gli3* | - | 3' | CRISPR/Cas9 | [2] |
| Human | *TEAD1* | 5' | 3' | Non-AUG start | [3] |
| Human | *MRVI1* | 5' | 3' | Non-AUG start | [3] |
| Human | *PIM1* | 5' | 3' | Non-AUG start | [3] |
| Human | *EIF4G2* | 5' | 3' | Non-AUG start | [3] |
| Human | *MYC* | - | 3' | Non-AUG start | [3] |
| Human | *RARB* | - | 3' | Non-AUG start | [3] |
| Human | *TEAD4* | 5' | 3' | Non-AUG start | [3] |
| Human | *R3HCC1* | 5' | 3' | Non-AUG start | [3] |
| Mouse | *Ctnnb1* | 5' | 3' | CRISPR/Cas9 | [4] |
| Human | *LMNA* | 5' | 3' | CRISPR/Cas9 | [4] |
| Zebrafish | *cd36* | 5' | 3' | ENU | [5] |
| Zebrafish | *pla2g12b* | 5' | - | ENU | [5] |
| Zebrafish | *smyd1* | 5' | - | CRISPR/Cas9 | [5] |
| Human | *PHACTR1* | 5' | 3' | CRISPR/Cas9 | [6] |
| Human | *CTNNB1* | 5' | 3' | CRISPR/Cas9 | [7] |
| Human | *AXIN1* | 5' | 3' | CRISPR/Cas9 | [7] |
| Human | *LRP5* | 5' | - | CRISPR/Cas9 | [7] |
| Human | *LRP6* | 5' | 3' | CRISPR/Cas9 | [7] |
| Human | *PTEN* | 5' | - | CRISPR/Cas9 | [7] |
| Human | *TBK1* | - | 3' | CRISPR/Cas9 | [7] |
| Human | *BAP1* | 5' | - | CRISPR/Cas9 | [7] |
| Human | *TLE3* | 5' | 3' | CRISPR/Cas9 | [7] |
| Human | *PPM1A* | - | 3' | CRISPR/Cas9 | [7] |
| Human | *BCL2L2* | 5' | 3' | CRISPR/Cas9 | [7] |
| Human | *NGLY1* | 5' | 3' | CRISPR/Cas9 | [8] |
| Human | *PI4KB* | 5' | 3' | CRISPR/Cas9 | [8] |
| Human | *BRD4* | 5' | 3' | CRISPR/Cas9 | [8] |
| Human | *BRD3* | - | 3' | CRISPR/Cas9 | [8] |
| Human | *MTF2* | 5' | 3' | CRISPR/Cas9 | [8] |
| Human | *BAZ2A* | 5' | 3' | CRISPR/Cas9 | [8] |
