## Supplemental Figures for "Genes adapt to outsmart gene targeting strategies in mutant mouse strains by skipping exons to reinitiate transcription and translation"

**Supplementary Figure 1.**

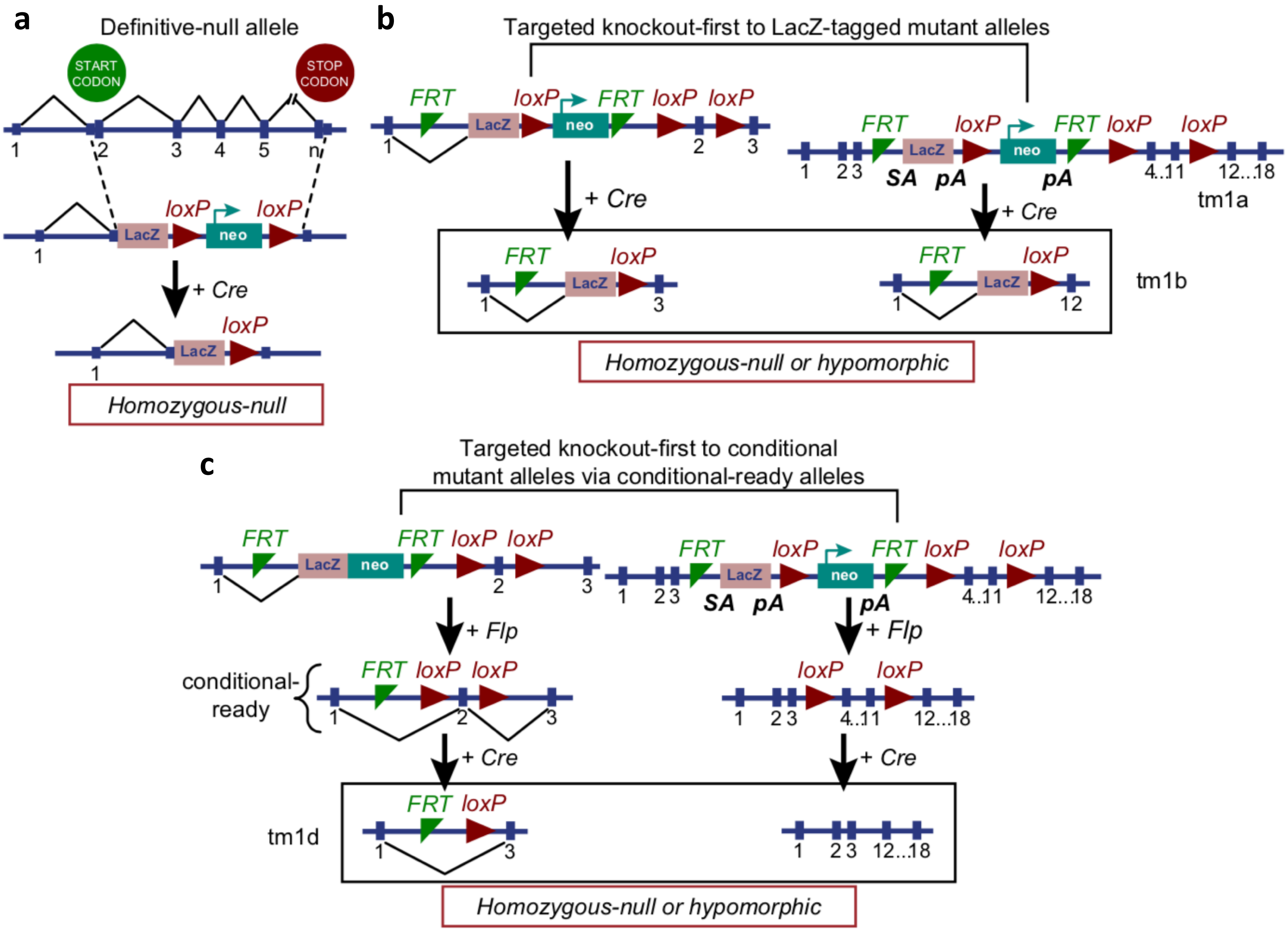

**Supplementary Figure 2.**

**a**

| | *Rhbdf1*$^{+/+}$ *Rhbdf2*$^{-/-}$ | *Rhbdf1*$^{+/-}$ *Rhbdf2*$^{-/-}$ | *Rhbdf1*$^{-/-}$ *Rhbdf2*$^{-/-}$ |
|---|---|---|---|
| | Number (%) Expected 25% | Number (%) Expected 50% | Number (%) Expected 25% |
| e9.5 | 4 (29%) | 8 (57%) | **2 (14%)** |
| e10.5 | 1 (4%) | 18 (75%) | **5 (21%)** |
| e12.5 | 5 (42%) | 6 (50%) | **1 (8%)** |
| e13.5 | 8 (40%) | 9 (45%) | **3 (15%)** |
| e14.5 | 4 (44%) | 4 (44%) | **1 (11%)** |
| e15.5 | 2 (25%) | 1 (12.5%) | **5 (62.5%)** |
| e17.5 | 2 (33%) | 2 (33%) | **2 (33%)** |
| e18.5 | 5 (33%) | 9 (60%) | **1 (7%)** |
| P1-3 | 6 (23%) | 16 (61%) | **4 (15%)** |
| >P4 | 9 (24%) | 29 (76%) | **0** |

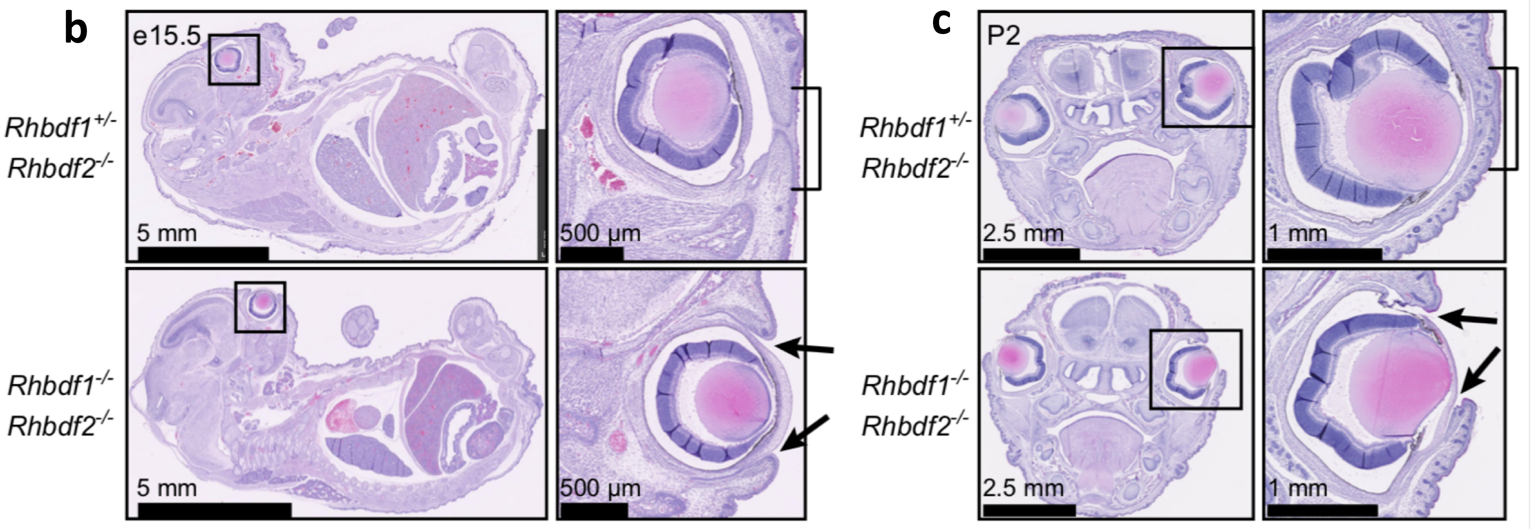

### Supplementary Figure 3

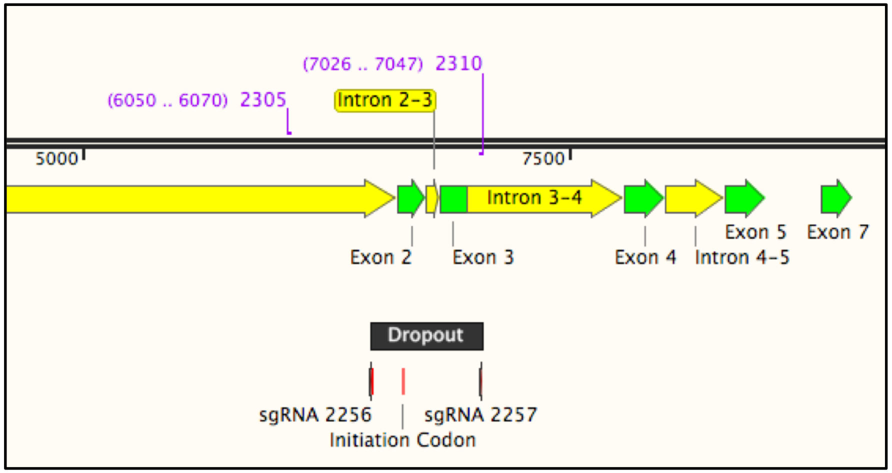


| Phusion High-Fidelity PCR | | | |
|---|---|---|---|
| CYCLE STEP | CYCLES | TEMP | TIME |
| Initial denaturation | 1 | 98°C | 30 seconds |
| Denaturation | | 98°C | 10 seconds |
| Annealing | 30 | 72°C | 30 seconds |
| Extension | | 72°C | 1 minute |
| Final extension | 1 | 72°C | 5 minutes |
| Hold | 1 | 4°C | ∞ |

| | |
|---|---|
| 2256 sgRNA | GGAGGATCTGTGGAGTTC |
| 2257 sgRNA | GAATCCCTGTTAGGTACC |

| | |
|---|---|
| 2305 Forward | AAGGAGAGTCCTCTGCATACT |
| 2310 Reverse | ACCTAACAGGGATTCTGTATGC |

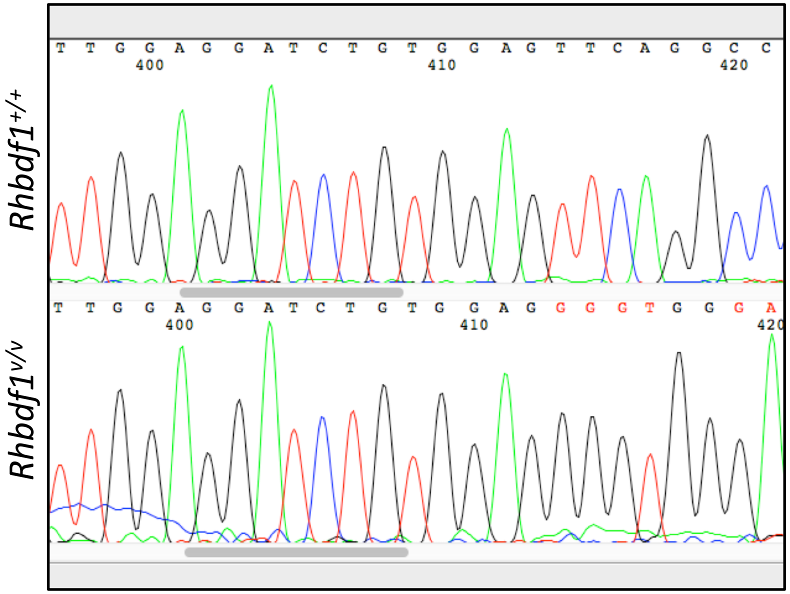


| | |
|---|---|
| Wildtype sequence | TTGTTGGAGGATCTGTGGAGTTCAGGCCCAGAGCATGCCA |
| Mutant sequence | TTGTTGGAGGATCTGTGGAG-GGGTGGGAATGGGGCAAAGG |

### Supplementary Figure 4

**a**

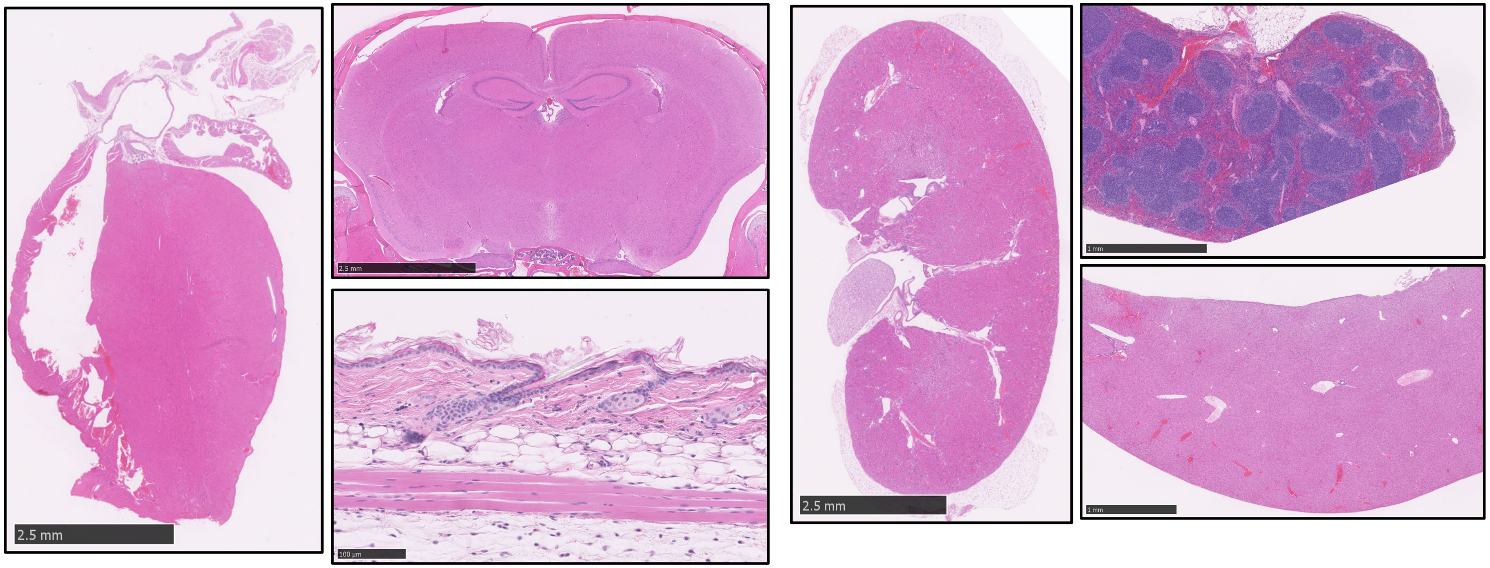


**b** Splice variants of *Rhbdf1*$^{v/v}$

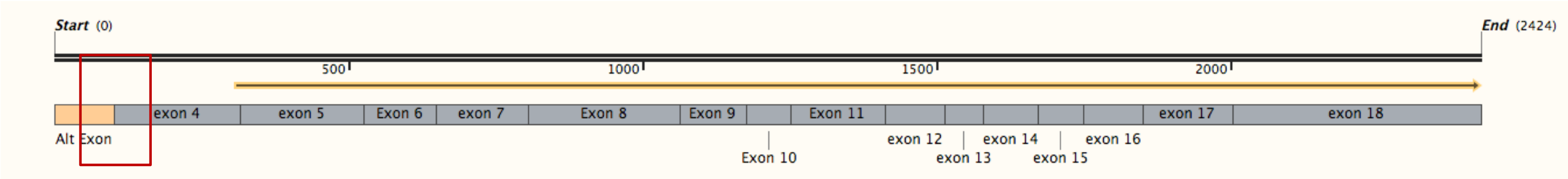


CAGGTACGCTTTGGGCGTGTTCATACTCTGCCCCTTTTGAGGCCGAGGGCTGCTCGAAAGGCCCACCCACAGCGCCAGTCTCTCTCATGGTCCTTGCTCAGGG
GGACAGCAGACTGGTTTGGAGTGAGCAAGGACAGTGACAGCACCCAGAAGTGGCAGCGCAAGAGCATCCGTCACTGCAGCCAGCGGTATGGGAAGCTAAA
GCCGCAGGTCATCCGGGAGCTGGACCTGCCCAGCCAG

| Transcript variant | NCBI Reference Sequence |
|---|---|
| X1 | XM_011243658.2 |
| X2 | XM_006514494.4 |
| X3 | XM_006514495.4 |
| X4 | XM_006514496.3 |
| X9 | XR_872202.2 |
| X14 | XR_001779873.2 |

### Supplementary Figure 5

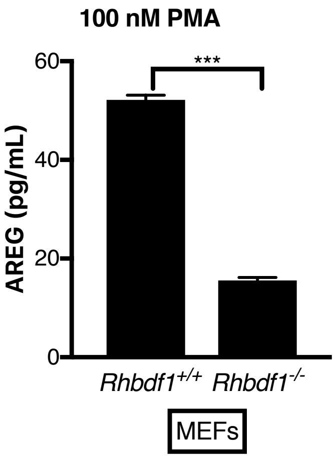

### Supplementary Figure 6

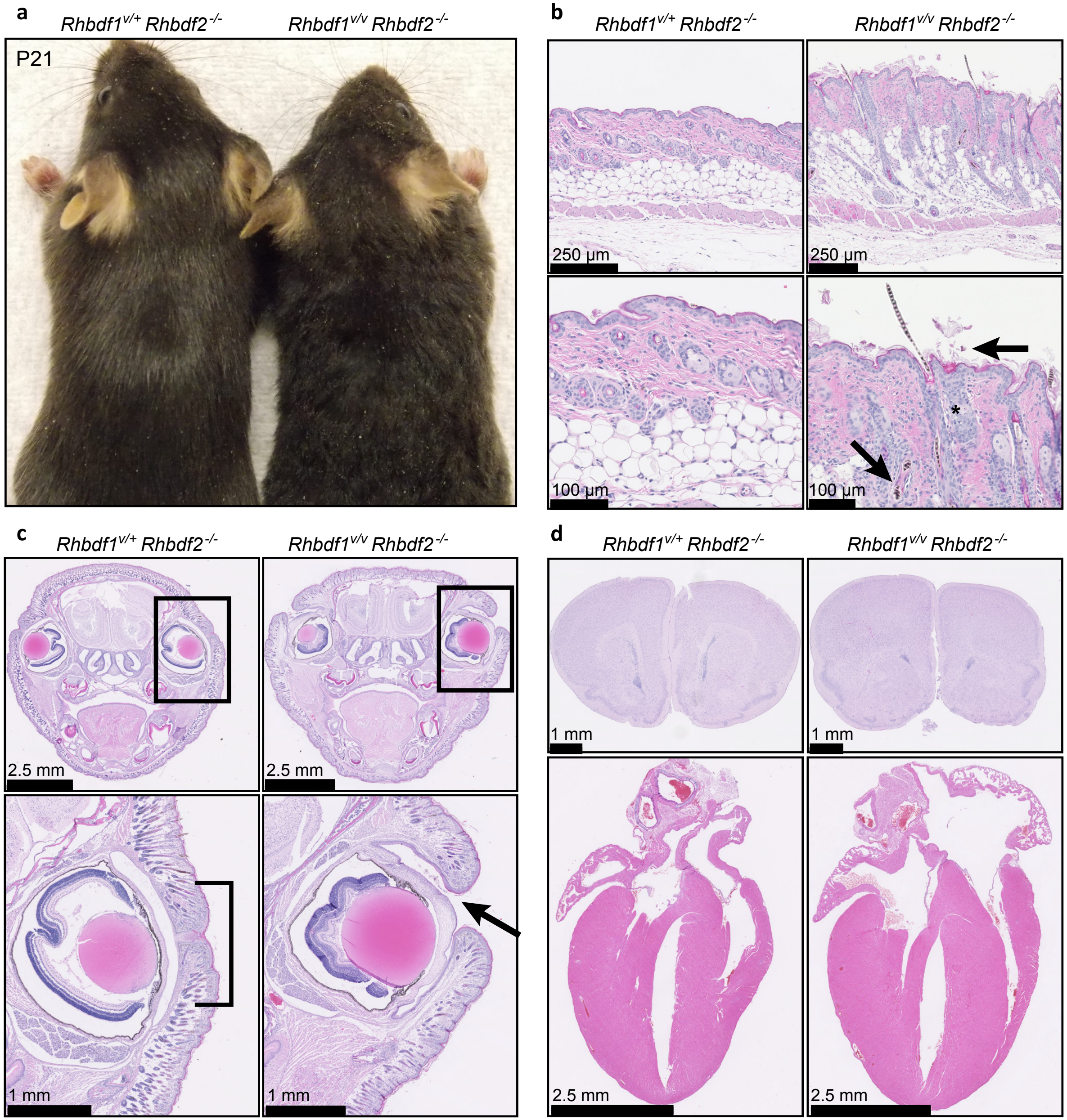

### Supplementary Figure 7

**a**

**Transcript: Mouse *Rhbdf1*-206 ENSMUST00000143988**

RPCCIGTKGRCEITSREYCDFMRGYFHEEATLCSQVHCMDDVCGLLPFLNPEMTVLRDLEKLAGWHRIAIIYLLSGITGNLASAIFLPYRAEVGPAGSQFGILACLFVELFQ
SWQILARPWRAFFKLLAVVLFLFAFGLLPWIDNFAHISGFVSGLFLSFAFLPYISFGKFDLYRKRCQIIIFQVVFLGLLAGLVVLFYFYPVRCEWCEFLTCIPFTDKFCEKYELD
AQLH*

**Transcript: Human *RHBDF1*-205 ENST00000448893**

RPCCIGTKGRCEITSREYCDFMRGYFHEEATLCSQVHCMDDVCGLLPFLNPEMTVLRDLEKLAGWHRIAIIYLLSGVTGNLASAIFLPYRAEVGPAGSQFGILACLFVELFQ
SWQILARPWRAFFKLLAVVLFLFTFGLLPWIDNFAHISGFISGLFLSFAFLPYISFGKFDLYRKRCQIIIFQVVFLGLLAGLVVLFYVYPVRCEWCEFLTCIPFTDKFCEKYELDA
QLH*

**b**

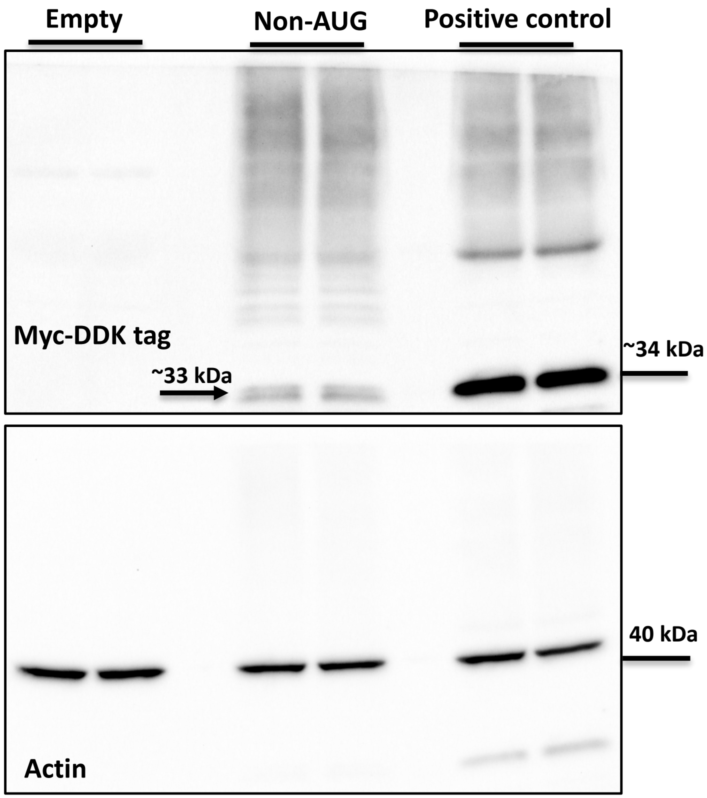

### Supplementary Figure 8

**a**

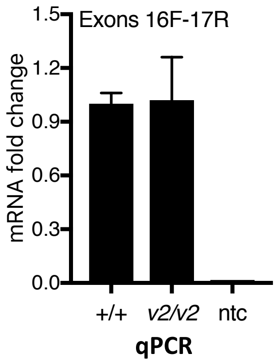


**b**

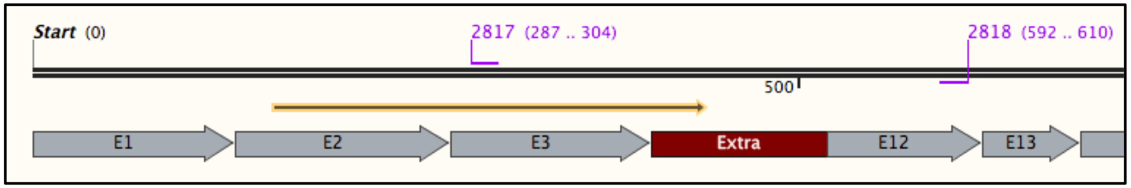


| | |
|---|---|
| Exon 3 2817 For | AGGCTTTCTTGCGGAGTG |
| Exon 12 2818 Rev | GGTGGCAGACAGAACCAAA |

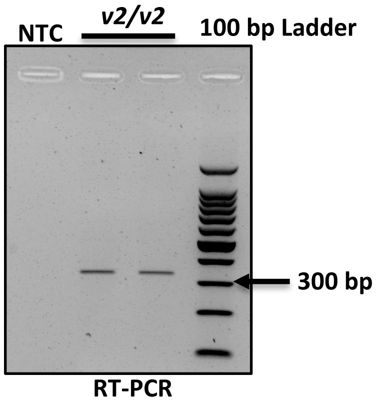


**c**

***AGGCTTTCTTGCGGAGTG***TGAGTATGCCAGCTGAGACAGCCCGGGTCCCATCACCCCACCATGAGCCTCGCCGGCTGGTGCT
GCAGCGTCAGACATCCATCACACAGACCATCCGCAGGTCCCAGGTCCCGAAAACCAAAGAAGAAGAACCCTAACAAAGAGG
ACAAGCGGCCTCGCACAGCCTTCACTGCTGAGCAGCTCCAGAGGCTCAAGGCTGAGTTTCAGACCAACAGTCCACGTTGGC
GGTGTGGGTGAAGTGGCCCGTTCATCCTAGTGCCCCAGACCTTGCAGGCAACAAGAGACAG***TTTGGTTCTGTCTGCCACC***

***Bold: Primers 2817/2818***

Extra: *En2* splice acceptor sequence from the targeted KO-first cassette

**d**

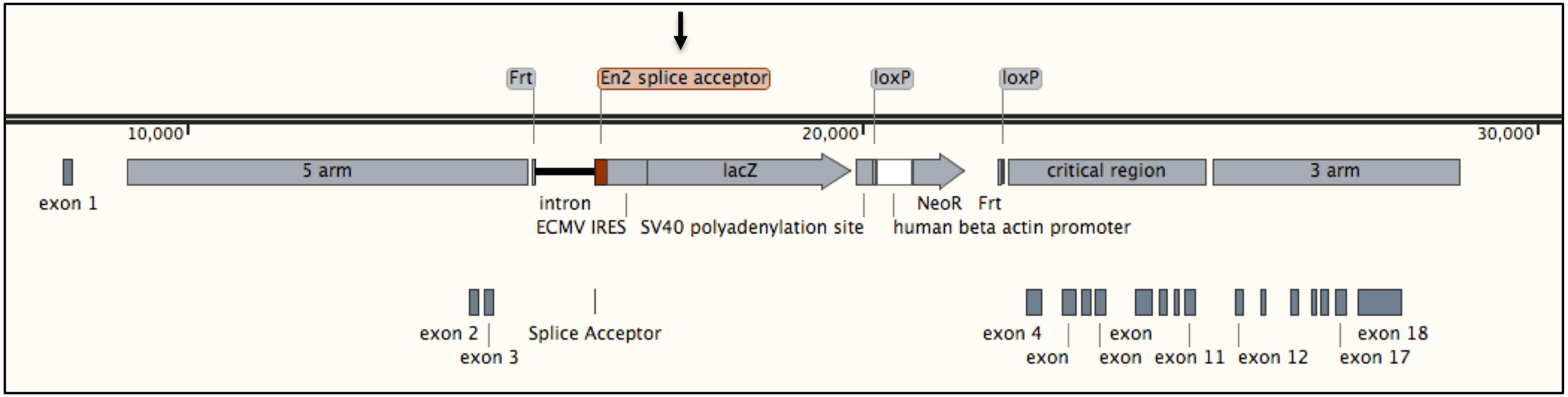


**e**

Putative 10.8 kDa truncated protein sequence:
MSEARRDSTSSLQRKKPPWLKLDIPAAVPPAAEEPSFLQPLRRQAFLRSVSMPAETARVPSPHHEPRRLVLQRQTSITQTIRRSQVPKTKEEEP*

**Supplementary Figure 9**

**a**

*Rhbdf1 targeted KO-first allele*

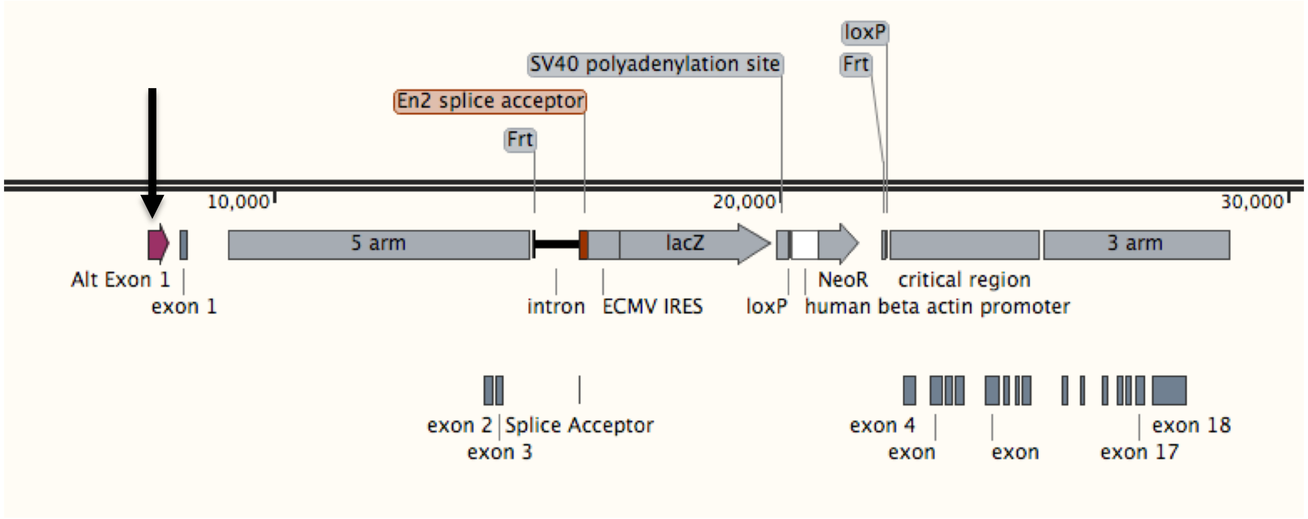


**b**

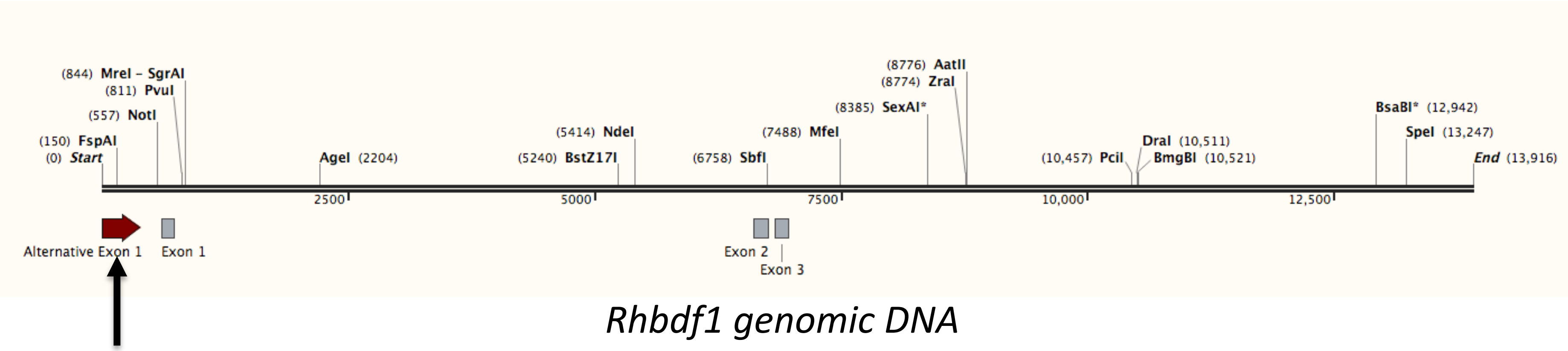


*Rhbdf1 genomic DNA*

<u>Alternative exon 1 sequence:</u>

tgtacctgctgggaaagaagtaggggaaggggtaattgctgagctgcggttcctgtgaaatggtgtatcagtgtatgaacgtgagtgtgcacacttataaatgtgttagggtagag
agtgtgagcccatggaagtgcattttgaataggtgtgcgcacgtgtctgtgtaaacaggtgcgagaggcctgaatgcatgaatgtttgttagaaagatcaggcgcgtgtgcacagg
acaagtgagagcttcgggaccctcagtgatgtgccagtgcgtgccttcggaaggggcgttcaatctgccttttaggtcccttcccacgccaccctcttgctttggaggtccccttcctc
tcttcccaagcacaggggcagcacaagatgaaggagtatggaaaggcgagtcca

### Supplementary Figure 10

**a**

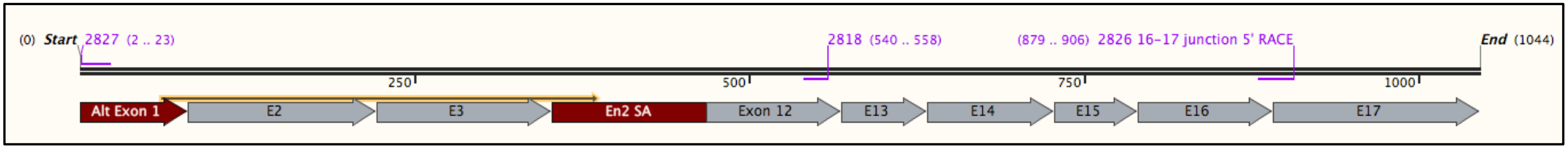


Putative 12.3 kDa truncated protein sequence:
MVYQCMNACLTAPGTMSEARRDSTSSLQRKKPPWLKLDIPAAVPPAAEEPSFLQPLRRQAFLRSVSMPAETARVPSPHHEPRRLVLQRQTSITQTIRRSQVPKTKEEEP*

**b**

| | |
|---|---|
| 2827 F | GTACCTGCTGGGAAAGAAGTAG |
| 2818 R | GGTGGCAGACAGAACCAAA |
| 2826 5' RACE | GATTACGCCAAGCTT CAAGCAGTGCAGGATCCCAGCGTGCAGG |

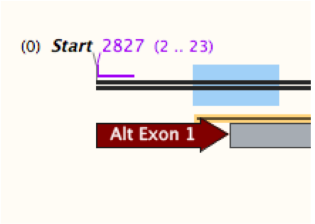


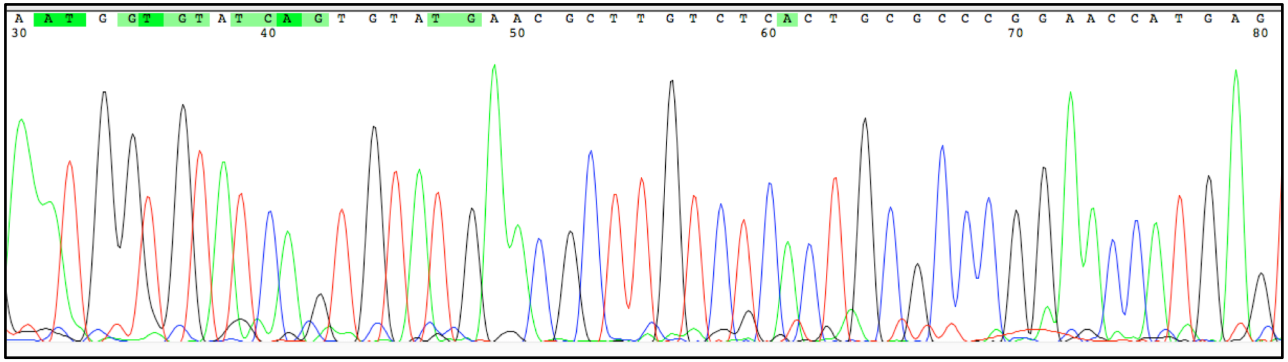

AATGGTGTATCAGTGTATGAACGCTTGTCTCACTGCGCCCGGAACCATGAG

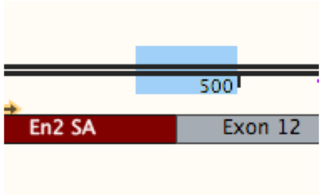


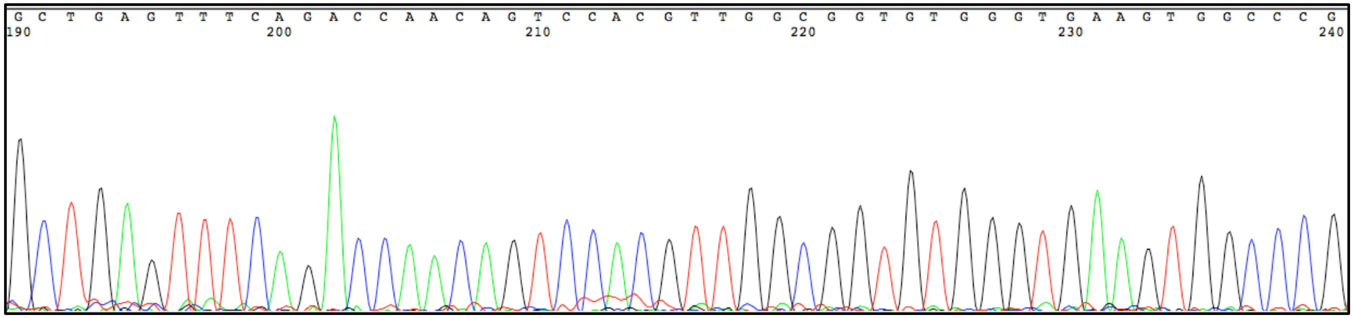

GCTGAGTTTCAGACCAACAGTCCACGTTGGCGGTGTGGGTGAAGTGGCCCG

### Supplementary Figure 11

**a**

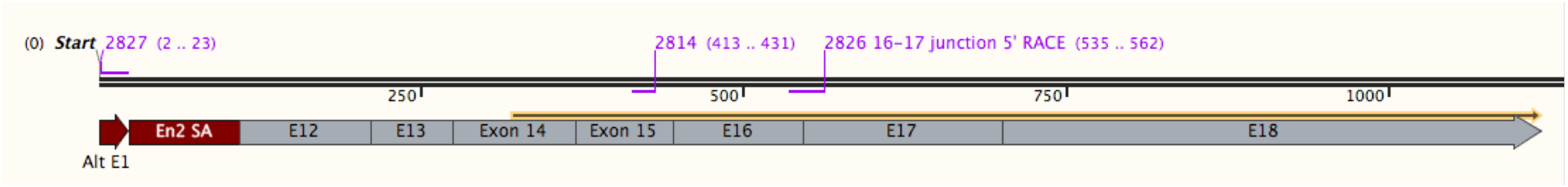


Variant 1; 32.77 kDa

MDCVITGRPCCIGTKGRCEITSREYCDFMRGYFHEEATLCSQVHCMDDVCGLLPFLNPEVPDQFYRLWLSLFLHAGILHCLVSVCFQMTVLRDLEKLAGWHRIAIIYLLSG
ITGNLASAIFLPYRAEVGPAGSQFGILACLFVELFQSWQILARPWRAFFKLLAVVLFLFAFGLLPWIDNFAHISGFVSGLFLSFAFLPYISFGKFDLYRKRCQIIIFQVVFLGLLA
GLVVLFYFYPVRCEWCEFLTCIPFTDKFCEKYELDAQLH(EQKLISEEDLDYKDDDDK)

**b**

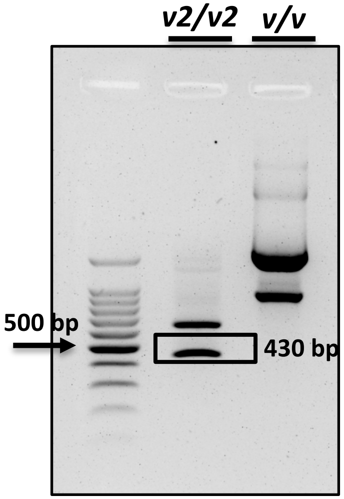


2827-2814 RT-PCR

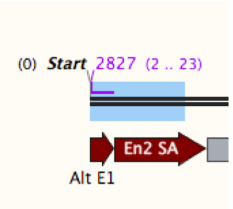


TGAGCCTCTGGAGCTGCTCAGCAGTGAAGGCTGTGCGAGGCCGCTT
GTCCTCTTTGTTAGGGTTCTACTTCTTTCCCAGCAGGTACA

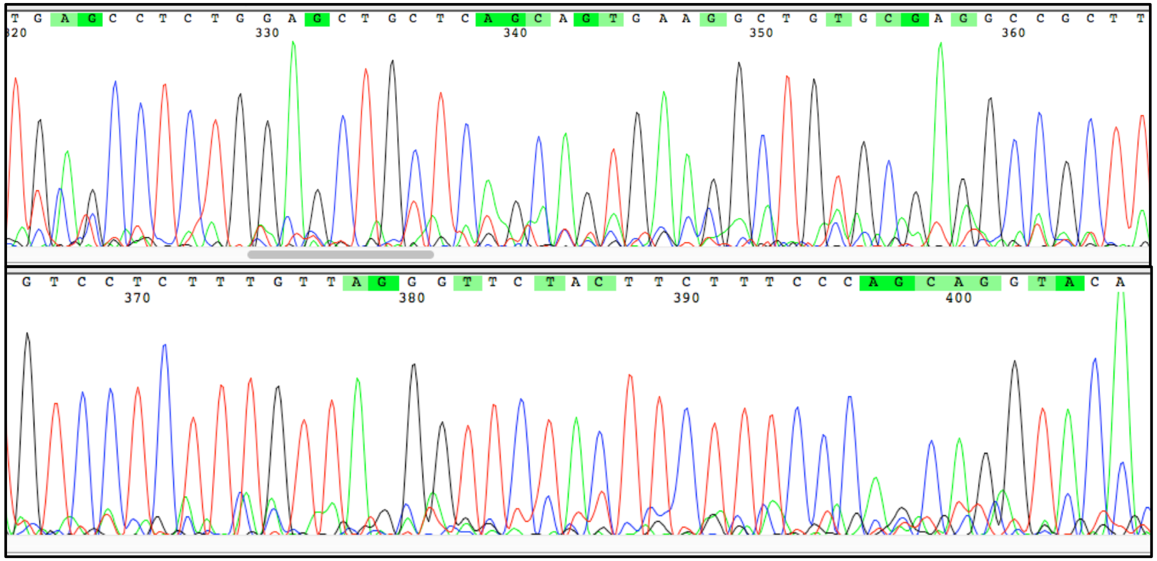

| Taq polymerase PCR | | | |
|---|---|---|---|
| CYCLE STEP | CYCLES | TEMP | TIME |
| Initial denaturation | 1 | 95°C | 2 minutes |
| Denaturation | 30 | 95°C | 15 seconds |
| Annealing | | 52°C | 30 seconds |
| Extension | | 68°C | 2 minute |
| Final extension | 1 | 72°C | 5 minutes |
| Hold | 1 | 4°C | ∞ |

**Supplementary Figure 12**

**a**

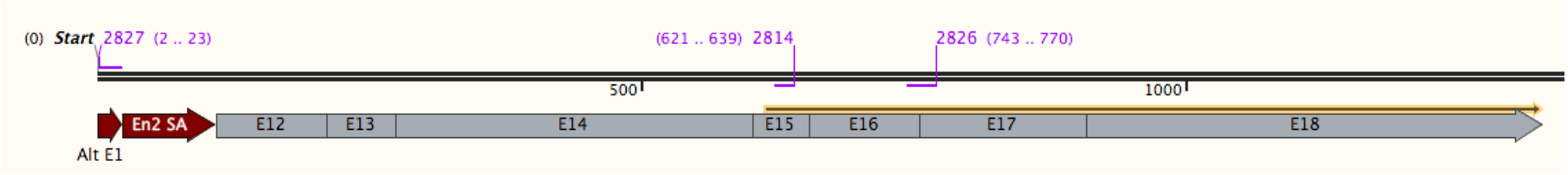


Variant 2; 29.6 kDa

MRGYFHEEATLCSQVHCMDDVCGLLPFLNPEVPDQFYRLWLSLFLHAGILHCLVSVCFQMTVLRDLEKLAGWHRIAIIYLLSGITGNLASAIFLPYRAEVGPAG
SQFGILACLFVELFQSWQILARPWRAFFKLLAVVLFLFAFGLLPWIDNFAHISGFVSGLFLSFAFLPYISFGKFDLYRKRCQIIIFQVVFLGLLAGLVVLFYFYPVRCE
WCEFLTCIPFTDKFCEKYELDAQLH(EQKLISEEDLDYKDDDDK)

**b**

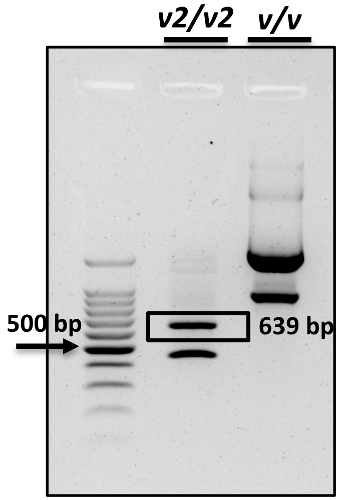


2827-2814 RT-PCR

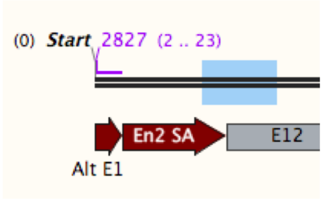


GCTGAGTTTCAGACCAACAGTCCACGTTGGCGGTGT
GGGTGAAGTGGCCCGTTCATCCTAG

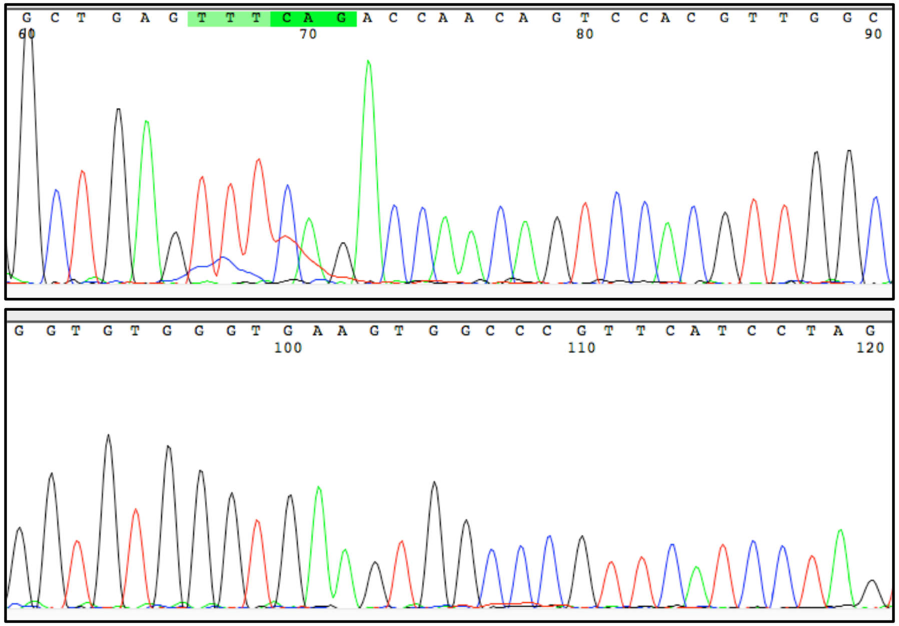

### Supplementary Figure 13

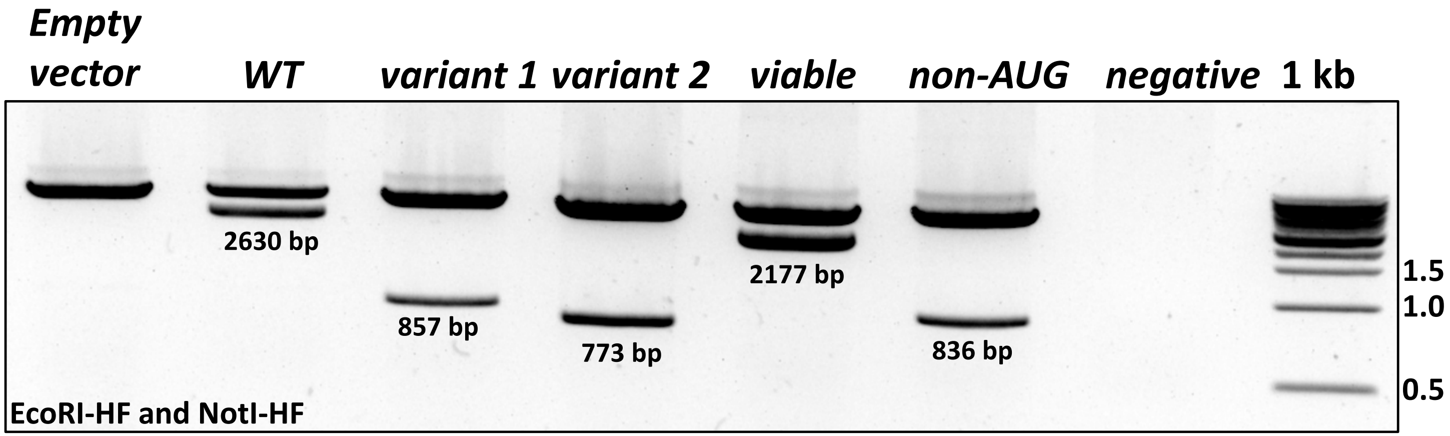

**Supplementary Figure 14**

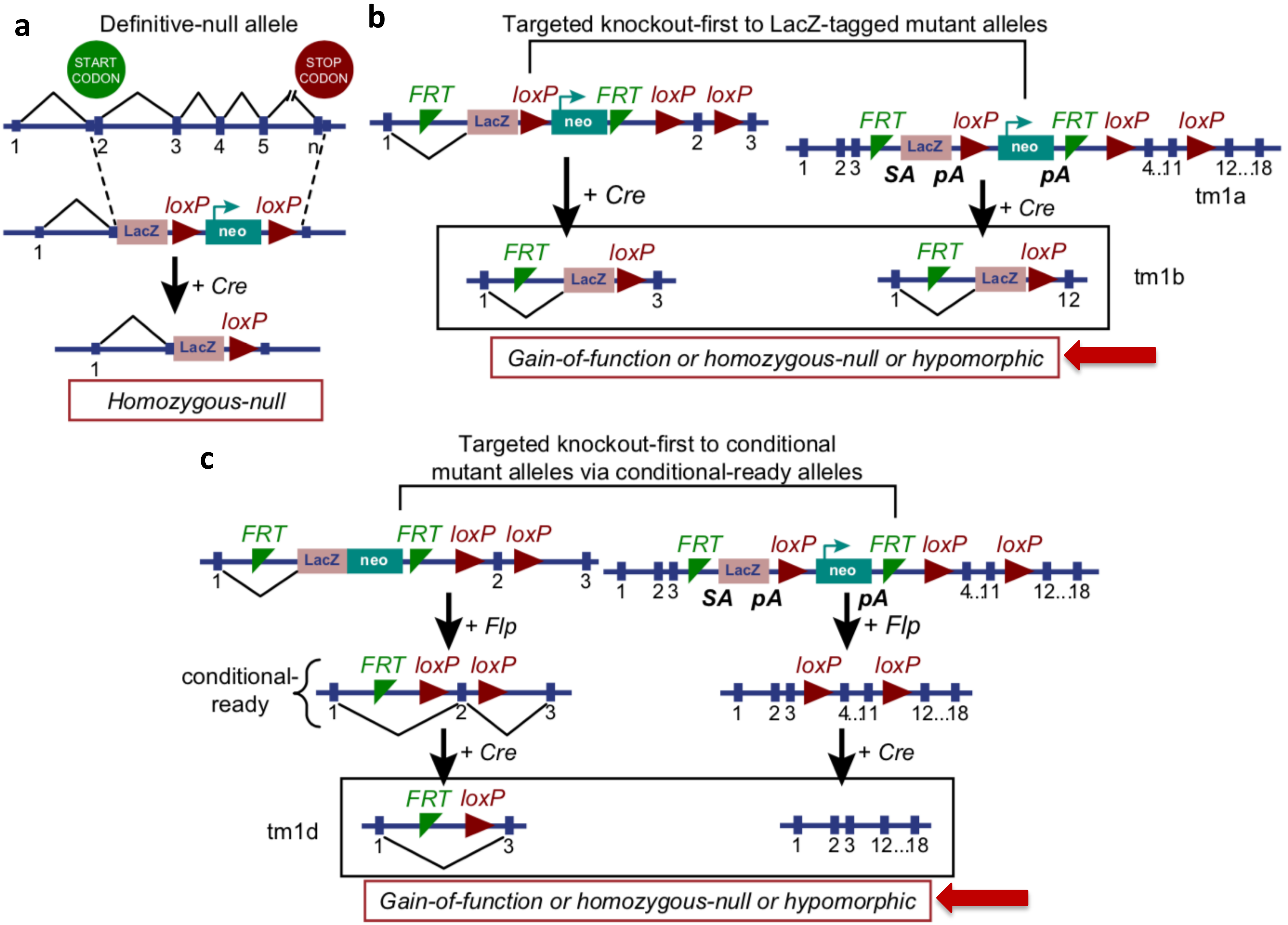

### Supplementary Figure 15

**a**

B6N(Cg)-*Rhbdf1*$^{tm1.1(KOMP)Vlcg}$/J

| | |
|---|---|
| Wt For | GCTCAGGTTTCAACCGACTC |
| Wt Rev | CAGGGAAGTCCACCATGTCT |
| Mt For | CGCTCCCAGTCCATTTATTG |
| Mt Rev | CGGTCGCTACCATTACCAGT |

| Taq polymerase PCR | | | |
|---|---|---|---|
| CYCLE STEP | CYCLES | TEMP | TIME |
| Initial denaturation | 1 | 95°C | 2 minutes |
| Denaturation | 30 | 95°C | 15 seconds |
| Annealing | | 62.5°C | 30 seconds |
| Extension | | 68°C | 2 minute |
| Final extension | 1 | 72°C | 5 minutes |
| Hold | 1 | 4°C | ∞ |

**b**

*Rhbdf2* KOMP allele

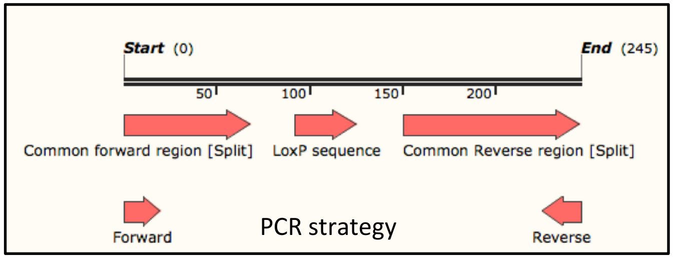


| | |
|---|---|
| Forward | AGTAGCATCTCCCCGTGGAT |
| Reverse | AACCAATGGACCATGGACCC |

| Taq polymerase PCR | | | |
|---|---|---|---|
| CYCLE STEP | CYCLES | TEMP | TIME |
| Initial denaturation | 1 | 95°C | 2 minutes |
| Denaturation | 30 | 95°C | 15 seconds |
| Annealing | | 55°C | 30 seconds |
| Extension | | 68°C | 2 minute |
| Final extension | 1 | 72°C | 5 minutes |
| Hold | 1 | 4°C | ∞ |

Wildtype

Heterozygous

Homozygous

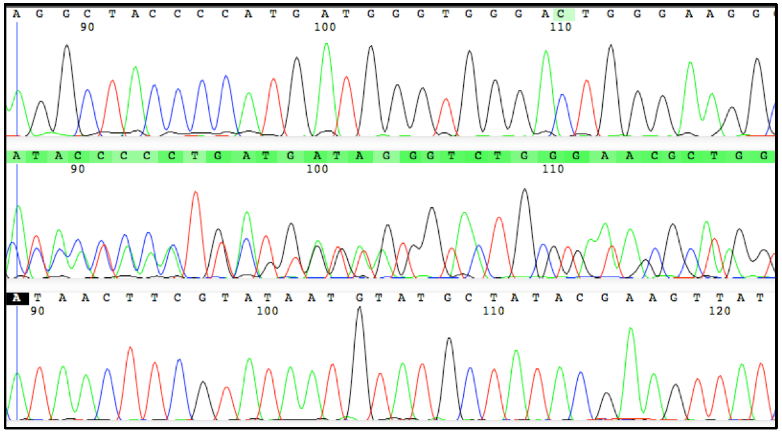
