## Supplemental Figure legends for "Genes adapt to outsmart gene targeting strategies in mutant mouse strains by skipping exons to reinitiate transcription and translation"

**Supplementary Figure 1. Gene-targeting strategies for generating null or conditional mutant alleles.**

a. In the definitive-null design, the complete coding sequence of a gene is flanked by two *loxP* sites, and recombination with a cre recombinase excises the floxed gene segment, creating LacZ-tagged null alleles. Loss of the complete coding sequence of the target gene ensures homozygous-null alleles.
b. Targeted KO-first alleles harbor *FRT* sites flanking "*LacZ* and a *neo* gene," and also harbor *loxP* sites flanking either a critical exon (exon 2, left) or specific exons chosen as critical (exons 4 through 11, right, in this case), permitting recombination with Flp or cre recombinases and enabling conditional mutagenesis. Application of cre recombinase deletes critical exons flanked by two *loxP* sites, generating LacZ-tagged null alleles (tm1b). Application of Flp recombinase to this allele is described in the legend for 1C.
c. Application of Flp recombinase to targeted KO-first alleles reverts the knockout-first alleles into conditional-ready alleles by excising *lacZ* and a *neo* gene. Subsequent recombination of conditional-ready alleles with cre recombinase generates conditional mutant alleles (tm1d).

**Supplementary Figure 2. Sub-viability in *Rhbdf1*$_{-/-}$ *Rhbdf2*$_{-/-}$ double mutant mice (A6)**

a. Breeding strategy and the number of mice generated and genotyped by PCR

b & c. *Rhbdf1*$_{-/-}$*Rhbdf2*$_{-/-}$ double mutant mice displayed "eyelids open at birth" phenotype (arrows) in mouse embryos harvested on embryonic day 15 (e15.5) (B) and in pups at day 2 (C). No defects were observed in *Rhbdf1*$_{+/-}$*Rhbdf2*$_{-/-}$ littermates.

**Supplementary Figure 3. PCR and genotyping of *Rhbdf1*$_{v/v}$ allele**

Schematic showing the TRU-guides (sgRNAs 2256 and 2257) used to excise exons 2 and 3 in the *Rhbdf1* gene. Also shown are the PCR primers and *Taq* DNA Polymerase thermocycling conditions used to genotype *Rhbdf1*$_{v/v}$ mice (A7). PCR products were sequenced to verify the 569-bp deletion.

**Supplementary Figure 4. Histology and 5' RACE products of *Rhbdf1*$_{v/v}$ allele**

a. H&E sections of *Rhbdf1*$_{v/v}$ mice displaying no abnormalities in the heart, brain, truncal skin sections, kidney, spleen, and liver tissues.
b. Schematic representation of *Rhbdf1*$_{v/v}$ transcript variant with a different transcription start site. 5' RACE identified an alternative exon (highlighted in light orange) that splices with exon 4 to reinitiate transcription from a different start site in exon 4, as opposed to using the conventional start site in exon 2. Notably, splicing of alternative exon with exon 4 is also observed in other *Rhbdf1* predicted transcript variants X1, X2, X3, X4, X9, and X14.

**Supplementary Figure 5.** Stimulated secretion of AREG in *Rhbdf1*$_{+/+}$ and *Rhbdf1*$_{-/-}$ primary MEFs, passage one, after overnight stimulation with 100 nM PMA. Following stimulation, cell-culture supernatants were analyzed using a mouse AREG ELISA kit. Data represent mean ± S.D; ***$p<0.001$.

**Supplementary Figure 6. CRISPR/Cas9 *Rhbdf1*$_{v/v}$ allele (A7) reverses sub-viability in *Rhbdf1*$_{-/-}$/*Rhbdf2*$_{-/-}$ *double mutant mice***

a. Images of representative postnatal day 21 *Rhbdf1*$_{v/+}$ *Rhbdf2*$_{-/-}$ mice (left) and *Rhbdf1*$_{v/v}$ *Rhbdf2*$_{-/-}$ mice (right). Notably, *Rhbdf1*$_{v/v}$ *Rhbdf2*$_{-/-}$ mice develop a wavy hair coat (arrow).
b. H&E-stained sections of skin from adult *Rhbdf1*$_{v/+}$ *Rhbdf2*$_{-/-}$ and *Rhbdf1*$_{v/v}$ *Rhbdf2*$_{-/-}$ mice. The *Rhbdf1*$_{v/v}$ *Rhbdf2*$_{-/-}$ skin displays follicular dystrophy (arrow), hyperplasia (*), and hyperkeratosis (arrowhead), whereas that of *Rhbdf1*$_{v/+}$ *Rhbdf2*$_{-/-}$ mice does not display these phenotypes. Top panels: low magnification; bottom panels: high magnification.
c. *Rhbdf1*$_{v/v}$ *Rhbdf2*$_{-/-}$ double mutant mice displayed an "eyelids open at birth" phenotype in 3-day-old mouse pups (arrow). No such defects were observed in *Rhbdf1*$_{v/+}$ *Rhbdf2*$_{-/-}$ mice. Top panels: low magnification; bottom panels: high magnification.
d. No cardiac or brain abnormalities were observed in either *Rhbdf1*$_{v/v}$ *Rhbdf2*$_{-/-}$ double mutant mice or *Rhbdf1*$_{v/+}$ *Rhbdf2*$_{-/-}$ mice.

**Supplementary Figure 7. Non-AUG protein coding transcripts of *Rhbdf1.***

a. Mouse non-AUG protein-coding *Rhbdf1* transcript (Top). Human non-AUG protein-coding *RHBDF1* transcript.
b. C-terminal Myc-DDK-tagged empty vector (lanes 1, 2), *non-AUG* vector (lanes 3, 4) or a positive control vector (798-bp; lanes 5, 6) were transiently expressed in 293T cells, and cell lysates were analyzed using western blotting with FLAG-specific antibody. Blots were washed, blocked in 5% nonfat dry milk, and re-probed with anti-actin antibody.

**Supplementary Figure 8. Retention of *En2* splice acceptor site in the *Rhbdf1*$_{v2/v2}$ allele following recombination with Flp and cre recombinases**

a. mRNA expression of the *Rhbdf1* gene assessed by RT-qPCR using SYBR Green RT-PCR master mix and, exon 16 forward and exon 17 reverse primers. There is no difference in the mRNA levels of *Rhbdf1* between *Rhbdf1*$_{+/+}$ and *Rhbdf1*$_{v2/v2}$ mice.
b. Schematic representation of *Rhbdf1*$_{v2/v2}$ allele indicating the presence of *En2* splice acceptor site (Extra) between exons 3 and 12. RT-PCR of spleens from *Rhbdf1*$_{v2/v2}$ mutant mice using primers in exon 3 and exon 12. Exon 3 forward (2817 For) and exon 12 reverse (2818 Rev) primers generated a 324-bp product, instead of an expected 209-bp product due to splicing of exons 3 and 12 together.
c. Sanger's sequencing indicated the presence of *En2* splice acceptor site (Extra) flanked by exons 3 and 12.
d. Schematic representation of the targeted KO-first allele used to generate ES cell clones displaying the splice acceptor site downstream of exon 3.
e. Transcription and translation from the conventional start site in exon 2 can yield a 10.8 kDa truncated protein owing to a stop codon in the splice acceptor site.

**Supplementary Figure 9. 5' RACE identifies alternative exon upstream of the conventional exon 1**

a. Schematic representation of the *Rhbdf1* KO-first allele used to generate ES cell clones displaying an alternative exon 1 (arrow) upstream of the conventional exon 1.
b. Schematic representation of the *Rhbdf1* genomic DNA displaying the alternative exon 1 (arrow) identified by 5' RACE. The DNA sequence of the alternative exon 1 is also shown.

**Supplementary Figure 10. Translation initiation from alternative exon 1 in the *Rhbdf1*$_{v2/v2}$ allele could generate a truncated protein**

a. Schematic representation of an *Rhbdf1*$_{v2/v2}$ splice variant displaying transcription initiation from the alternative exon 1, which could result in a 285-bp transcript and a 12.3 kDa truncated protein.
b. Primers (2827, 2818, and 2826) used to sequence verify splicing of alternative exon 1 with exon 2 (top), and *En2* splice acceptor site (*En2* SA) with exon 12 (bottom).

**Supplementary Figure 11. Splicing of alternative exon 1 with exon 4 generates an N-terminally truncated variant 1 protein**

a. Schematic representation of an *Rhbdf1*$_{v2/v2}$ splice variant displaying transcription initiation from the exon 14, which could result in a 801-bp transcript and a 32.77 kDa N-terminally truncated protein (variant 1).
a. RT-PCR of spleens from *Rhbdf1*$_{v2/v2}$ mutant mice using primers (2827 and 2814) in alternative exon 1 and exon 15 resulted in a 430-bp amplicon. Sanger's sequence demonstrated splicing of alternative exon 1 with *En2* SA. Notably, *Rhbdf1*$_{v/v}$ mutant mice resulted in an amplicon with primers 2827 and 2814, suggesting the use of alternative exon 1. However, higher molecular weight PCR fragments in *Rhbdf1*$_{v/v}$ mutant mice can be attributed to the presence of exons 4 through 11, which are excised in *Rhbdf1*$_{v2/v2}$ mutant mice. Thermocycling conditions using *Taq* DNA Polymerase with standard buffer are also shown.

**Supplementary Figure 12. Splicing of alternative exon 1 with exon 4 can also generate an N-terminally truncated variant 2 protein**

a. Schematic representation of an *Rhbdf1*$_{v2/v2}$ splice variant displaying transcription initiation from the exon 15, which could result in a 717-bp splice variant and a 29.6 kDa N-terminally truncated protein (variant 2).
b. RT-PCR of spleens from *Rhbdf1*$_{v2/v2}$ mutant mice using primers (2827 and 2814) in alternative exon 1 and exon 15 also resulted in a 639-bp amplicon. Sanger's sequence demonstrated splicing of *En2* SA with exon 12.

**Supplementary Figure 13.** Double digests of C-terminal Myc-DDK-tagged vectors (1 μg DNA) with NEB's restriction enzymes EcoRI-HF and NotI-HF.

**Supplementary Figure 14.** Our data strongly suggests that mice harboring homozygous definitive-null or homozygous targeted KO-first alleles are homozygous null, however, mice harboring homozygous KO-first mutant alleles tm1b and tm1d can be null or gain-of-function or hypomorphic mice (arrows) owing to unpredicted reinitiation of transcription and translation.

**Supplementary Figure 15. PCR and genotyping of *Rhbdf1* and *Rhbdf2* KOMP alleles**

a. PCR primers used to genotype *Rhbdf1*$^{-/-}$ mice (A5) generated in the present study. Thermocycling conditions using *Taq* DNA Polymerase with standard buffer.
b. Schematic showing the PCR strategy and the primers used to genotype *Rhbdf2*$^{-/-}$ mice. Thermocycling conditions using *Taq* DNA Polymerase with standard buffer. PCR products were purified and sequence verified.
